# Photolabile oligonucleotides combined with topologically imposed light gradients enable spatially resolved single-cell transcriptomics and epigenomics

**DOI:** 10.1101/2025.10.01.679703

**Authors:** Robert A. Piscopio, Alex Chialastri, Chieh Wang, Mikolaj Godzik, Kellie A. Heom, Weiyue Wang, Maxwell Z. Wilson, Siddharth S. Dey

**Affiliations:** Department of Molecular, Cellular, and Developmental Biology, University of California Santa Barbara, Santa Barbara, CA 93106, USA; Department of Chemical Engineering, University of California Santa Barbara, Santa Barbara, CA 93106, USA; Department of Biological Engineering, University of California Santa Barbara, Santa Barbara, CA 93106, USA; Neuroscience Research Institute, University of California Santa Barbara, Santa Barbara, CA 93106, USA

## Abstract

The organization of cells within a tissue plays a critical role in tuning cellular function. Several methods have recently been developed to capture the transcriptome of cells while retaining spatial information. However, these genome-wide sequencing methods typically lack the spatial resolution of individual cells and are confined to quantifying positional information within predefined lattice locations, thereby failing to capture large sections of a tissue outside these regions. Further, these methods are generally limited to profiling fixed cells with reduced mRNA capture efficiency compared to standard scRNA-seq. In addition, existing methods lack modularity and cross-platform compatibility, thereby limiting most of these techniques from jointly profiling the epigenetic and transcriptomic state of individual cells. To overcome these limitations, we present scSTAMP-seq (<u>s</u>ingle-<u>c</u>ell <u>S</u>patial <u>T</u>ranscriptomic <u>A</u>nd <u>M</u>ultiomic <u>P</u>rofiling), an approach that employs cholesterol-tagged photolabile oligonucleotides that incorporate into cell membranes, enabling us to “stamp” the position of cells using spatially imposed light gradients prior to tissue dissociation and single-cell sequencing. Applied to live cells, scSTAMP-seq efficiently captures spatially resolved single-cell transcriptomes at high resolution for all cells within a field of view. Further, we demonstrate that light patterning enables dynamic spatial resolution, including the ability to map the position of individual cells. Finally, we show that scSTAMP-seq is modular and can be seamlessly integrated with various downstream single-cell sequencing technologies. We demonstrate this by performing scRNA-seq using plate- and droplet-based methods, and by performing joint epigenome and transcriptome sequencing from the same cell while preserving positional information. Collectively, these results demonstrate that scSTAMP-seq is a sensitive and high-throughput technology for mapping single-cell transcriptomes and epigenomes at the spatial resolution of individual cells.

## INTRODUCTION

Tissue function relies on the coordinated action of cell types across multiple length scales^1–4^. For example, juxtacrine interactions between neighboring cells are coupled with longer range paracrine signaling to determine spatiotemporal patterns of gene expression at the single cell level. Modern single-cell RNA sequencing (scRNA-seq) methods are optimized for high-throughput collection of single-cell transcriptomes from heterogenous tissues; however, these platforms necessitate the dissociation of a sample into individual cells, which results in a loss of positional information and renders the investigation of spatial coordination among cell types impossible^5,6^. The same challenges exist with single-cell epigenome sequencing, thereby limiting our understanding of how the spatial structure of a tissue integrates with epigenetic reprogramming to control transcriptional programs of individual cells. Therefore, we sought to develop a modular method that leverages the advantages of existing single-cell sequencing platforms to simultaneously profile the epigenome and transcriptome of individual cells while preserving spatial information by dynamically etching spatial coordinates onto a sample using projected light and photocleavable oligonucleotides embedded in cell membranes.

Broadly, two modalities of spatial transcriptomics have emerged. The first modality uses iterative on-microscope optical quantification of transcripts that can achieve single-cell and subcellular spatial resolution with high transcript detection accuracy. However, these methods are limited to fixed samples that make it unfeasible to capture tissue dynamics, employ a pre-defined set of probes that limit the scale of the transcriptome captured, and generally cannot quantify the epigenetic landscape of individual cells^7–16^. The second modality, grid- or spot-based sequencing methods, impart a spatial dimension onto samples using DNA barcoded arrays, beads or microwells arranged in a lattice; however, they suffer from a variety of limitations result from the grid or spot patterns^1–3,17–27^. Fixed lattices are generally larger than single cells, resulting in the transcriptome of multiple cells being conflated to a single spot. Additionally, these methods have “dead regions” between pixels to prevent cross-contamination of transcripts resulting in the inability to capture large sections of a tissue, and are constrained to profiling a fixed cross-sectional area of a tissue determined by the size of predesigned barcoded microarrays or microfluidic devices. Further, as these spots/grids are printed in a uniform pattern, these methods are unable to dynamically alter the spatial resolution within a tissue. Furthermore, these methods generally also have lower efficiency of capturing unique mRNA molecules compared to established scRNA-seq methods, possibly as the mRNA is captured on spots/grids from fixed cells. Another key limitation of using fixed samples is that these methods cannot capture morphogenesis and cell migration in real time. Finally, most spatial sequencing methods cannot quantify the epigenome or make joint single-cell epigenome and mRNA measurements. Therefore, an improved spatial sequencing method would ideally integrate the best of both modalities, enabling genome-wide quantification of the transcriptome and epigenome at a single-cell resolution. To accomplish this, such a method would (1) enable sample-specific scaling of pixel resolution as a function of tissue complexity, (2) be unconstrained by the manufactured size of any pixel bed without containing any dead regions, (3) be compatible with live cells, and (4) allow for both transcriptome and epigenome measurements to be made on single cells.

A promising third modality for spatial transcriptomics has emerged that uses light to encode spatial information. Two methods in particular, Zip-seq and Light-seq, have the potential to overcome some of the challenges encountered in the first two modalities by leveraging the precise, yet highly reconfigurable patterning of light onto samples. However, both methods use binary encoding (presence or absence of light) of positional information, thereby resulting in a more limited spatial resolution. Further, Light-seq profiles fixed samples, resulting in low transcript detection efficiency, limited depth of coverage, and the inability to quantify cell migration. Finally, neither method can capture the epigenome or make integrated measurements of both the epigenome and transcriptome from single cells. To overcome these challenges, we present scSTAMP-seq (<u>S</u>ingle-cell <u>S</u>patial <u>T</u>ranscriptomic <u>A</u>nd <u>M</u>ultiomic <u>P</u>rofiling), which improves upon existing light-based methods by utilizing a photolabile oligonucleotide that enables spatially resolved single-cell measurements of the transcriptome and epigenome from live cells. With user-defined projected light patterns that can be modulated in intensity (256 bit), duration (∼1 ms), and spatial precision (∼500 nm), we demonstrate that “stamping” cells can be used to gain insights into how spatial signals and the epigenome regulate specific gene expression programs in a cell.

## RESULTS

### PHO barcodes coupled with patterned light illumination achieves flexible spatially resolved single-cell transcriptome profiling

scSTAMP-seq utilizes photocleavable hashtag oligonucleotides (PHOs) that incorporate into cell membranes to optically embed positional information onto cells using spatially patterned light before a sample is dissociated and subjected to single-cell sequencing (Fig. 1a). scSTAMP-seq PHOs are designed with an ‘inner’ and ‘outer’ DNA sequence that is linked by a photocleavable moiety (Fig. 1a and Supplementary Fig. 1a). The ‘inner’ DNA sequence hybridizes to cholesterol-conjugated “anchor” oligonucleotides that enable PHO embedding into cell membranes (Fig. 1a). Thus, quantifying the extent of PHO cleavage, together with mRNA from the same cell, enables spatially resolved single-cell transcriptomics using scSTAMP-seq. While not required to spatially encode cell location into a sequencing readout, the ‘inner’ and ‘outer’ sequence of the PHOs can be conjugated with distinct fluorophores allowing the extent of PHO cleavage to be visualized prior to sequencing (Supplementary Fig 1b and Supplementary Table 1). These additional indicators also enable enrichment for cells of interest at specific spatial locations via cell sorting. Finally, both PHO ‘inner’ and ‘outer’ DNA sequences contain a handle for downstream PCR amplification, an ‘inner’- and ‘outer’-specific barcode, a random 6-nucleotide unique molecule identifier (UMI) to count individual molecules, and a poly-A capture sequence (Supplementary Fig. 1a,c).

**Figure 1.**
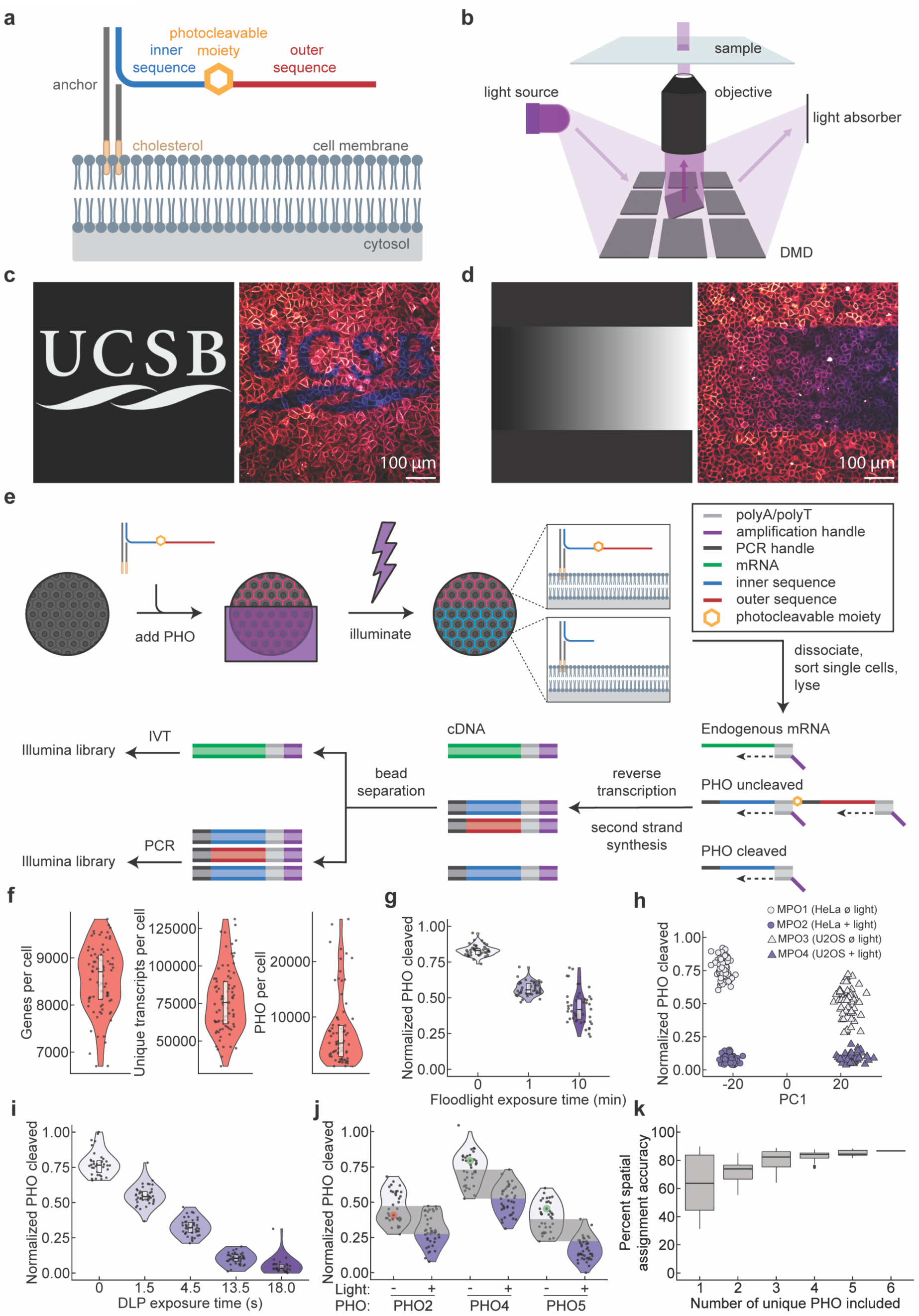
scSTAMP-seq enables spatially-resolved single-cell transcriptomics by employing photo-sensitive molecules and light to encode positional information. **a**, Schematic of photocleavable hashtag oligonucleotide (PHO) and cholesterol modified oligonucleotide (CMO) scaffolds used to label the membrane of live cells. **b**, Schematic of DMD orientation and function for precise user-defined light patterning. **c**, Image of a binary mask to illuminate cells using a DMD (left). False-color composite image of overlapping fluorescence from ‘inner’ sequence (blue) and ‘outer’ sequence (magenta) of PHOs post-illumination with a binary mask (right). **d**, Image of a linear light gradient mask (left) applied to cells and the resulting false-color composite fluorescence image after illumination (right). **e**, Workflow of scSTAMP-seq. The first step involves embedding PHO molecules in the cell membrane of live cells via CMO scaffolds. A light mask is then applied to photocleave the ‘outer’ sequence (red) of PHOs. Samples are then dissociated, sorted as single cells, and lysed. Reverse transcription followed by second strand synthesis is performed on endogenous mRNA and PHOs. Next, the PHO-derived molecules are separated from mRNA-derived molecules using a paramagnetic bead small molecule enrichment. Endogenous mRNA and PHO libraries are then prepared separately, where the mRNA-derived molecules are amplified via *in vitro* transcription (IVT) and subsequent PCR, while the PHO-derived molecules are amplified by PCR to generate final Illumina libraries. **f**, Violin plots of the number of genes detected per cell, the number of transcripts detected per cell, and the number of PHO molecules detected per cell using plate-based cell capture and sequencing in scSTAMP-seq. Dots represent individual cells. **g**, Violin plots of ‘outer’-to-‘inner’ read counts of PHOs in each cell captured using plate-based sequencing for cells exposed to three distinct illumination exposure times. **h**, Cell-type classification of HeLa and U2OS cell lines using transcriptome-based principal component analysis, ‘outer’-to-‘inner’ PHO read counts and MPOs. **i**, Violin plot of ‘outer’- to-‘inner’ PHO read counts in each cell for a uniquely barcoded PHO after 5 distinct illumination exposure times. Dots represent individual cells. **j**, Violin plot demonstrates error correction when multiple uniquely barcoded PHOs are used simultaneously in the same experiment. Highlighted dots (red and green) represent the same cell, where red indicates ambiguous light-dose assignment and green indicates corrected light-dose assignment. **k**, Bar plots show the accuracy of spatial assignment of single cells as a function of uniquely barcoded PHOs included in the analysis.

We first sought to understand the degree to which PHO cleavage is light-dose dependent. In scSTAMP-seq, samples are exposed to user-defined light patterns by employing digital light processing (DLP) projection via a digital micromirror device (DMD) to direct a 360-380 nm LED light source (Fig. 1b). As an initial proof-of-concept to demonstrate that the combination of light patterning and PHOs can be used to spatially label cells, we applied a UCSB logo photomask to stamp over a live cell monolayer incubated with PHOs containing Cy3 and Cy5 fluorophores, conjugated to the ‘inner’ and ‘outer’ DNA sequences, respectively (Fig. 1c, Supplementary Fig. 1b, 2a, and Supplementary Video 1). As a separate demonstration to show that light can be used beyond a binary readout of cellular location, a linear light gradient was applied to assess the extent to which the cleavage fraction of PHOs could be spatially controlled. We found that the cleaved fraction of PHOs in each cell is monotonically correlated to light exposure (Fig. 1d, Supplementary Fig. 2b,c and Supplementary Video 2). This suggests that spatially guided differential light exposure can be used to molecularly encode cellular position based on the fraction of cleaved PHOs. In addition to DLP, we also used a 365 nm LED floodlight as a convenient alternative to illuminate a larger sample area to show that both illumination methods can successfully cleave PHOs (Supplementary Fig. 2c and 3a,b). Finally, to demonstrate that this method has single-cell spatial precision in a complex monolayer, we co-cultured U2OS cells with HeLa cells expressing mCherry. We showed that by specifically targeting the mCherry positive HeLa cells with user-defined light patterns, we could cleave the PHOs from these cells at high specificity compared to neighboring U2OS cells (Supplementary Fig. 4a,b). Overall, these experiments demonstrate that custom light gradients, together with PHOs, can be used to spatially labels individual cells in scSTAMP-seq.

Next, to establish scSTAMP-seq as a spatially resolved single-cell transcriptomics technology, we developed the following method. First, PHOs are incubated with samples to embed these molecules into cell membranes (Fig. 1e). Next, samples are illuminated with user-defined light patterns, prior to dissociation and single-cell sorting into individual wells of a 384- well plate (Fig. 1e). After cell lysis, a poly-T primer with an overhang containing a cell-specific barcode, a 6-nucleotide UMI, a 5’ Illumina adapter and a T7 promoter is used to reverse transcribe endogenous mRNA molecules and copy the ‘inner’ and ‘outer’ DNA sequences of the PHOs (Supplementary Fig. 1c). Thereafter, second-strand synthesis is performed and the individual wells from a 384-well plate are pooled as the mRNA molecules and PHOs are tagged with cell-specific barcodes (Supplementary Fig. 1c). Paramagnetic solid phase reversible immobilization (SPRI) beads are then used to separate the short PHO products from the mRNA-derived cDNA (Supplementary Fig. 1c, 5). The PHO-derived molecules are then amplified by PCR to generate Illumina libraries, and the mRNA-derived cDNA molecules are amplified by *in vitro* transcription (IVT) followed by IIlumina library preparation, as described previously^28–33^. Thus, simultaneous sequencing of PHOs and cDNA from the same cell enables spatially resolved single-cell transcriptomics in scSTAMP-seq.

As proof-of-concept, we applied scSTAMP-seq to cells exposed to three distinct levels of illumination to find that we can successfully quantify both PHOs and the transcriptome from single cells (Fig. 1f). Compared to scRNA-seq, we detected a similar number of unique transcripts and genes per cell in scSTAMP-seq, suggesting that the quality of the transcriptome obtained from scSTAMP-seq is equivalent to standard scRNA-seq (Supplementary Fig. 6). Importantly, we found that the fraction of PHOs cleaved in individual cells is directly proportional to light exposure, showing that the spatial location and corresponding light dosage can be directly linked to sequencing output, enabling spatially resolved single-cell transcriptomics (Fig. 1g and Supplementary Fig. 3a,b). Finally, as the spatial position of a cell is inferred from the fraction of PHOs cleaved (ratio of detected ‘outer’ to ‘inner’ DNA sequences of PHOs), this measurement is robust to noise arising from incorporation efficiency of the PHOs into cell membranes.

We next tested the general applicability of scSTAMP-seq. First, we determined that the UV exposure required for PHO cleavage does not alter the endogenous transcriptome of these cells, confirming that light exposure in scSTAMP-seq does not induce transcriptional artifacts (Supplementary Fig. 7a,b). Next, to demonstrate that PHOs are not incorrectly assigned to cells in scSTAMP-seq due to the transfer of these molecules between cells after sample dissociation and/or during cell sorting, two different cell lines, HeLa and U2OS, were incubated with multiplex oligos (MPOs) as described previously^28,29^, and exposed to distinct illumination levels to show that individual cells could be accurately identified based on their transcriptome, MPOs, or the fraction of PHOs cleaved, confirming that there is no cross-contamination PHOs between single cells (Fig. 1h and Supplementary Fig. 1a). Overall, these results demonstrate that PHOs and user-defined light gradients, combined with downstream single-cell sequencing, can be used to obtain spatially resolved single-cell transcriptomes in scSTAMP-seq.

### PHO multiplexing achieves noise reduction and increases spatial resolution

While our initial experiments showed that the spatial position of cells and the extent of light exposure is directly proportional to the fraction of cleaved PHOs, at each light exposure level we also observed a spread in the fraction of cleaved PHOs across a population of cells (Fig. 1g). We hypothesized that this uncertainty arises from technical variability in the measurement of ‘inner’ and ‘outer’ DNA sequence counts of the PHOs associated with each cell, thereby limiting the spatial resolution of the method. If the spread arises from random processes rather than systematic biases, then the simultaneous use of multiple uniquely barcoded PHOs should behave as an error correction feature by reducing measurement noise. Therefore, we incubated cells with six uniquely barcoded PHOs and exposed them to five distinct light exposure levels. Analysis of the six unique PHOs showed that individually, each displayed the expected monotonic relationship of increased PHO cleavage with increasing light exposure (Fig. 1i and Supplementary Fig. 8a,b). However, the PHO cleavage fraction for a small number of cells fell within the overlapping region of two adjacent distributions, implying that in the absence of prior knowledge, determining the light levels these cells were exposed to and their corresponding spatial location would not be possible. However, by incorporating the cleavage fraction from multiple unique PHOs on the same cell (see Methods for details), we could correct for this ambiguity and accurately assign the light exposure and spatial location of individual cells (Fig. 1j). Quantitatively, we found that inclusion of each additional PHO increased the accuracy with which we could assign the spatial location of individual cells, with the utilization of all six unique PHOs enabling >85% accuracy in cell assignment (Fig. 1k). In summary, these results demonstrate that incorporation of multiple uniquely barcoded PHOs decreases technical variability and increases the spatial resolution of scSTAMP-seq.

We hypothesized that we could further increase the spatial resolution of scSTAMP-seq by employing sequential rounds of optical labeling with uniquely barcoded PHOs. To demonstrate this, we used a combinatorial barcoding approach to spatially segment four quadrants of a cell monolayer (Fig. 2a). Briefly, after incubation with a unique PHO (PHO1), quadrants Q2, Q3, and Q4 were exposed to saturating levels of light to ensure complete cleavage of the PHOs in these regions (Fig. 2a,b and Supplementary Fig. 9a – step 1). Thereafter, we added PHO2 and exposed Q3 and Q4 again to saturating illumination (Supplementary Fig. 9a – step 2). Lastly, we added PHO3 and illuminated Q4 with saturating light (Supplementary Fig. 9a – step 3). This strategy ensures that each quadrant has a unique combination of cleaved and uncleaved PHOs (Fig. 2b). For visualization of this patterned PHO labeling, the ‘outer’ sequences of PHO1-3 were conjugated with distinct fluorophores (Fig. 2c, and Supplementary Fig. 1b, 9a,b). To connect the transcriptome to the spatial location of individual cells within one of the four quadrants, we subsequently processed these colonies using scSTAMP-seq to find that by quantifying the cleavage fraction of all three PHOs, we could assign the spatial location of cells with high precision (Fig. 2d and Supplementary Fig. 9b). Thus, this combinatorial barcoding approach demonstrates that iterative cycles of PHO labeling and custom light illumination patterns improves the spatial resolution of scSTAMP-seq.

**Figure 2.**
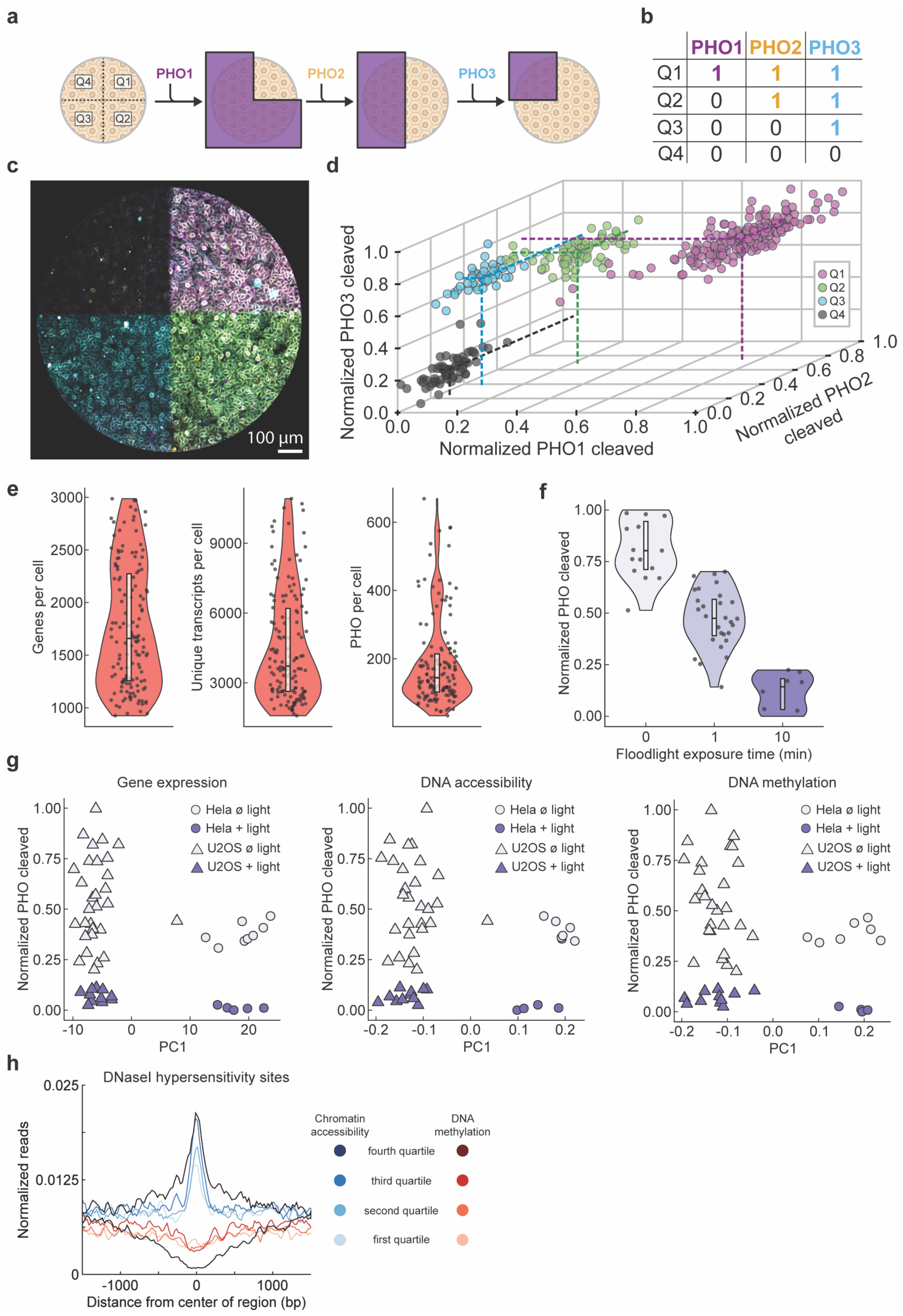
scSTAMP-seq is compatible with sequential rounds of optical labeling and its modularity enables spatially-resolved single-cell multiomics. **a**, Schematic shows that the sequential addition of uniquely barcoded PHOs and custom illuminations patterns allow cells within different spatial compartments to be labeled with a unique combination of cleaved and uncleaved PHOs. **b**, Table describes the cleavage status of PHO1-3 in the four distinct quadrants generated using three binary photomasks. ‘1’ indicates that a PHO is not cleaved and ‘0’ indicates that a PHO is cleaved. **c**, False-color fluorescence image of PHO1, PHO2 and PHO3 overlayed in magenta, yellow and cyan, respectively, over a cell monolayer. **d**, 3D plot of ‘outer’-to-‘inner’ PHO read counts for three uniquely barcoded PHOs on each cell. Individual cells are colored corresponding to assigned quadrant based on the combination of ‘outer’-to-‘inner’ PHO read counts. Dotted lines correspond to the average cleavage for each uniquely barcoded PHO in each quadrant. **e**, Violin plots of the number of genes detected per cell, the number of transcripts detected per cell, and the number of PHO molecules detected per cell using 10X droplet-based cell capture and sequencing in scSTAMP-seq. **f**, Violin plots of ‘outer’-to-‘inner’ PHO read counts in each cell captured using 10X droplet-based sequencing for cells exposed to three distinct illumination exposure times. **g**, scSTAMP-MAT-seq shows that HeLa and U2OS cells exposed to saturating or no light can be classified based on the endogenous transcriptome, DNA accessibility or DNA methylation profiles. **h**, Averaged single-cell DNA accessibility (blue) and DNA methylation (red) profiles from scSTAMP-MAT-seq at DNase I hypersensitivity sites in HeLa cells, split by previously reported signal strength^35^.

### scSTAMP-seq is compatible with droplet-based single-cell transcriptomics and single-cell multiomics methods

A notable aspect of scSTAMP-seq is its inherent compatibility with other single-cell sequencing methods. This is achieved through the orthogonal nature of PHO embedding and light patterning used to encode spatial information, which is independent of downstream sample dissociation and sequencing. This modularity enables us to combine scSTAMP-seq with any single-cell sequencing technology that also captures poly-adenylated molecules. As an initial demonstration of this versatility, we extended the plate-based single-cell profiling presented above to droplet-based cell capture and sequencing with the scSTAMP-seq pipeline to enable spatially resolved single-cell transcriptomics. Similar to the experiment described in Figure 1f,g, cells were incubated with PHOs and then exposed to three distinct illumination levels, followed by downstream processing on a 10X platform (with minor modifications, see Methods for details) to successfully quantify both endogenous transcripts and the ‘inner’ and ‘outer’ sequences of PHOs (Fig. 2e and Supplementary Fig. 10). We again confirmed that light exposure does not alter the endogenous transcriptome, validating that combining scSTAMP-seq with other single-cell sequencing technologies does not introduce transcriptional artifacts (Supplementary Fig. 11a,b). Finally, as before, we found that the fraction of PHOs cleaved was directly proportional to light exposure, suggesting that scSTAMP-seq can be combined with 10X droplet-based profiling to quantify spatially resolved single-cell transcriptomes (Fig. 2f).

The modularity of scSTAMP-seq further provides an opportunity to spatially profile the epigenome and transcriptome from single cells, thereby providing deeper insights into how cell extrinsic tissue structure and cell intrinsic epigenetic landscapes regulate gene expression. We have recently developed several single-cell multiomics methods that can simultaneously quantify both the epigenome and the transcriptome from the same cell, including scMAT-seq, a method to jointly profile DNA methylation, DNA accessibility, and the transcriptome from single cells^30,31,34^. To encode spatial information into scMAT-seq measurements, we have extended the technology as follows (termed scSTAMP-MAT-seq). After labeling cells with PHOs and spatially imposed light gradients, single cells from dissociated samples are sorted into 384-well plates and the following two steps are performed simultaneously – (1) mRNA is reverse transcribed and the ‘inner’ and ‘outer’ sequences of PHOs are copied using a poly-T primer with an overhang containing a cell- and mRNA/PHO-specific barcode, a UMI, a 5’ Illumina adapter and a T7 promoter; and (2) a methyltransferase, M.CviPI, is used to methylate cytosines in a GpC context within open chromatin (Supplementary Fig. 12). Performing these two steps simultaneously minimizes mRNA degradation and ensures the *in vivo* state of chromatin is captured immediately after cell lysis. Next, after second strand synthesis, 5-hydroxymethycytosine (5hmC) sites in the genome are glucosylated to block downstream detection by the restriction enzyme MspJI. Thereafter, MspJI is added, which recognizes methylated cytosines in the genome and creates double-stranded (ds)-DNA breaks that are ligated to ds-adapters containing a cell- and genomic DNA (gDNA)- specific barcode, a UMI, a 5’ Illumina adapter and a T7 promoter. Following this, as all molecules are tagged with cell- and molecule-of-origin-specific barcodes, individual wells of the 384-well plate are pooled, and paramagnetic SPRI beads are used to separate the short PHOs from barcoded mRNA- and gDNA-derived molecules. As described in Figure 1e, the PHO-derived molecules are then amplified by PCR to generate Illumina libraries. The mRNA- and gDNA-derived molecules are amplified by IVT, followed by Illumina library preparation, as described previously^31^. Finally, depending on the context of the methylated cytosine in the sequencing data, gDNA reads are either assigned to the methylome (CpG context) or DNA accessibility (GpC context) dataset for each cell. Thus, by combining scSTAMP-seq with downstream molecular barcoding in scMAT-seq, scSTAMP-MAT-seq can quantify spatially resolved DNA methylation, DNA accessibility, and mRNA from the same cell.

We performed scSTAMP-MAT-seq on HeLa and U2OS cell lines where both lines were exposed to saturating- or no-light conditions. Compared to scRNA-seq and scSTAMP-seq, we found that the number of genes and unique transcripts detected per cell in scSTAMP-MAT-seq were similar (Supplementary Fig. 13). In addition, the number of endogenously methylated cytosines detected per cell, and the number of methylated GpC sites detected per cell (corresponding to DNA accessibility), were similar for both scMAT-seq and scSTAMP-MAT-seq (Supplementary Fig. 14). Further, we were able to distinguish the cells either based on their transcriptome, methylome or DNA accessibility profiles, as well as identify the spatial position of these cells based on the extent of PHO cleavage (Fig. 2g). Finally, comparison of scSTAMP- MAT-seq with previously published DNase I hypersensitivity sites showed that more accessible sites were associated with larger DNA accessibility peaks and greater domain spreading, as well as lower levels of DNA methylation (Fig. 2h)^35^. Overall, these results show that scSTAMP-seq is modular and can be used to perform integrated epigenome and transcriptome sequencing from the same cell while maintaining positional information.

## DISCUSSION

In this work, we present scSTAMP-seq, a technology that leverages photocleavable hashtag oligonucleotides, custom light illumination patterns, and single-cell sequencing to accurately track cellular position and quantify single-cell transcriptomes. The strength of scSTAMP-seq lies in its unique barcoding system, where light exposure initiates cleavage of the PHOs, thereby “stamping” the spatial location of each cell that can be visualized if desired and then further decoded through downstream sequencing. We demonstrate here how the granularity of this spatial information can be enhanced through multiplexed and sequential barcoding, which significantly reduces technical noise and improves spatial resolution. Finally, the modularity of scSTAMP-seq allows it to be seamlessly integrated with other technologies, such as plate- or droplet-based single-cell transcriptomics, as well as comprehensive multiomics profiling at a single-cell level. Our demonstration of scSTAMP-MAT-seq underscores this potential, showing that this method can quantify spatially resolved DNA methylation, DNA accessibility, and mRNA simultaneously from individual cells. This integrated data provides an unparalleled opportunity to correlate gene expression dynamics with reprogramming of the epigenetic landscape within the same cell, contextualized within its spatial coordinates.

In the future, we plan to address several potential limitations of scSTAMP-seq. First, the current PHO configuration limits cleavage to a single wavelength of light and does not retain “memory” of cleavage patterns so that future illumination events do not interfere with past impressions. Therefore, future versions of scSTAMP-seq may include additional photolabile chemistries and methods for making PHOs insensitive to light during subsequent illumination events of sequential barcoding rounds. Second, the accuracy of scSTAMP-seq in more complex tissues, particularly those with pronounced three-dimensional structures, remains to be explored.

Illumination devices in three dimensions could employ two-photon illumination methods or holography to etch cellular voxels, and methods for delivering PHOs into dense three-dimensional tissue need to be systematically explored.

Overall, scSTAMP-seq offers a versatile and modular platform for spatially resolved single-cell transcriptomics, together with convenient integration with other single-cell multiomics methods. Future applications of scSTAMP-seq can significantly advance our understanding of the interplay between cellular spatial organization and epigenome and transcriptome dynamics in dissecting complex biological systems and disease processes.

## METHODS

### scSTAMP-seq oligo design

#### Photocleavable Hashtag Oligo (PHO)

PHO molecules contain a 50-nucleotide ‘inner’ and 50- nucleotide ‘outer’ sequence that are linked by a photolabile moiety sensitive to light in the UV-spectrum (360-380 nm). The ‘inner’ and ‘outer’ DNA sequences were both designed to have 4 components: (1) a 20-nucleotide handle for downstream PCR amplification, (2) a 4-nucleotide ‘inner’- or ‘outer’-specific barcode, (3) a random 6-nucleotide unique molecule identifier (UMI) to count individual molecules, and (4) a 20-nucleotide poly-A capture sequence. The ‘inner’ PCR handle is also complementary to CMO anchors to embed PHOs in cell membranes. A schematic of PHO components can be found in Supplementary Figure 1a. The intensity and/or duration of illumination of each PHO determines the amount of the ‘outer’ barcode sequence relative to the unchanging amount of the ‘inner’ barcode sequence. This ‘outer’-to-‘inner’ ratio is the foundation for encoding spatial information using scSTAMP-seq. Photocleavage rate is measured through ‘outer’-to-‘inner’ barcode read counts, where ‘inner’ barcode counts inform total PHOs a cell received, and ‘outer’ barcode counts inform illumination exposure. For image-based quality control, quantification, and confirmation of photocleavage prior to downstream Illumina library preparation, Cy5, Cy3, and FAM fluorophores were conjugated to PHOs. We designed three different PHO fluorophore conjugation schemes: (i) the 3’ ends of both the ‘inner’ and ‘outer’ sequences, (ii) the 3’ end of the ‘outer’ sequence, or (iii) neither sequence (no visual readout). For scheme (i), we conjugated Cy3 to the ‘inner’ sequence and Cy5 to the ‘outer’ sequence. For scheme (ii), we conjugated Cy3, Cy5, and FAM to the ‘outer’ sequence only. Initial experiments were conducted with scheme (i). After initial proof-of-concept experiments verifying PHO embedding and photocleavage efficiency, we opted for schemes (ii) and (iii) for later experiments. A schematic of the different fluorophore conjugated PHO designs can be found in Supplementary Figure 1b.

#### Multiplex Oligo (MPO)

MPO molecules are similar in design to the ‘inner’ sequence of PHOs, but do not contain a photolabile moiety or ‘outer’ sequence. MPOs contain a 20-nucleotide handle for CMO binding and downstream PCR amplification, a 4-nucleotide multiplex ID for distinguishing MPOs from PHOs, a 6-nucleotide barcode for distinguishing between different MPOs, a 6- nucleotide UMI, and a 20-nucleotide poly-A capture sequence. MPOs allow samples with different conditions and variables to be pooled and identified downstream by demultiplexing multiplex barcode reads to increase sample processing efficiency. MPOs are amplified simultaneously with PHOs in library preparation. A schematic of MPO components can be found in Supplementary Figure 1a.

#### Cholesterol Modified Oligo (CMO)

CMO anchor and co-anchor were adapted from Weber *et al.*^28^ and McGinnis *et al.*^29^, with slight modifications to DNA sequences. CMO anchors contain a 5’ cholesterol-TEG group for PHO cell membrane embedding, a 20-nucleotide sequence that hybridizes to the ‘inner’ handle of PHO molecules, a 20-nucleotide sequence that hybridizes to CMO co-anchors, and a 3’ amino modifier to prevent unwanted CMO amplification in downstream library preparation steps. CMO co-anchors contain a 3’ cholesterol-TEG group and a 20- nucleotide sequence that hybridizes to CMO anchors.

Detailed sequences of all scSTAMP-seq oligo sequences can be found in Supplementary Table 1. PHO, MPO, and CMO sequences were designed by the authors and synthesized by Integrated DNA Technologies (IDT). All DNA modifications are commercially available from IDT.

### Oligonucleotide and barcode sequences

Supplementary Table 1

### Oligo annealing and stock preparation

#### Resuspending and storing oligos

PHOs, MPOs, and CMOs were resuspended in nuclease-free water (Invitrogen, AM9932) to create 100 µm primary stocks. Primary stocks were aliquoted, protected from light using amber Eppendorf tubes (Eppendorf, 022363221), and stored at -20°C to prevent multiple freeze-thaw cycles.

#### Annealing oligos

An equimolar solution containing 2.4 µL PHO (100µM), 2.4 µL CMO anchor (100µM), and 7.2 µL of duplex buffer (IDT, 11-05-01-03) was pipetted into PCR tubes (Corning, 3745). This 20 µM intermediate stock solution was then placed in a Bio-Rad thermal cycler and heated to 94°C for 3 minutes. The temperature was slowly decreased by 1°C per minute until 4°C was reached to allow for oligo hybridization to occur. The solution was then held at 4°C. A 500 nM working stock of this solution was created immediately before an experiment and then kept on ice or placed at -20°C in a light-proof container for later use. For MPO-CMO anchor annealing, the same hybridization strategy as described above was used with MPOs replaced for PHOs.

### Cell culture

HeLa and U2OS cells were cultured and maintained on tissue cultured treated plastic flasks (Thermo Scientific, 156367) or 6-well plates (Corning, 3516) in DMEM with high glucose, GlutaMAX and sodium pyruvate (Gibco, 10569044), supplemented with 10% fetal bovine serum (FBS) (Gibco, 10569044) and 1% Pen-Strep (Gibco, 15140122). All mammalian cell cultures were maintained in incubators at 37°C and 5% CO_2_. Cells were routinely passaged 1:6 after 70-80% confluence was reached using 0.25% trypsin-EDTA (Gibco, 25200056). For microscopy imaging, cells were seeded onto glass 96 well glass bottom plates (Cellvis, P96-1.5H-N) or 35 mm dishes with glass coverslips (MATTEK, P35G-1.5-20-C) coated with fibronectin (Advanced Biomatrix, 5050). For the combinatorial barcoding experiment, cells were seeded on a custom protein patterned 96-well plate (CYTOO, 20-950-00) treated with fibronectin.

### PHO labeling of cell membranes

First, cell culture media was removed, and cells were washed three times with cold DPBS+/+ (with calcium and magnesium) (Gibco, 14040133). Hybridized PHO/CMO-anchor at 500 nM was then incubated with live-cell samples over ice for 15 minutes. Next, cells were washed again three times with cold DPBS+/+. After washing, CMO co-anchors were incubated with live-cell samples over ice for 15 minutes. Cells were then washed again three times with cold DPBS+/+. This subsequent two-step anchor/co-anchor incubation is optimal for oligo embedding rate and retention time, as described previously by Weber *et al.*^28^. Cells were then imaged and illuminated with UV light. After illumination, cells were either processed for single cell sorting, or the labeling procedure was repeated for experiments that utilized iterative rounds of unique spatial barcode labeling. For experiments that incorporated MPOs, both PHOs and MPOs were incubated simultaneously.

### Microscopy and photomask illumination

#### Microscopy imaging setup

All microscopy was performed using a Nikon Ti-2 inverted confocal microscope equipped with a Yokogawa CSU-W1 SoRa spinning disk and an Andor iXon Life 888 EMCCD camera. Images were captured with a Nikon CFI S Plan Fluor ELWD 20x objective lens and a Nikon CFI Plan Apo Lambda 10x objective lens. For fluorescence confocal imaging, 488 nm, 561 nm, and 640 nm lasers at 100% power were delivered through liquid light guides for FAM, Cy3, and Cy5 fluorophores, respectively. Digital light processing (DLP) patterned light projection was conducted using a Mightex Polygon400-G digital micromirror device (DMD) coupled with an Excelitas X-Cite 360-380 nm UVA liquid light guide-coupled LED, or a 365 nm 50W LED floodlight (Everbeam, B08635F9CX) with an acrylic light diffusion film and custom 3D printed plate stand for consistent and uniform illumination.

#### UV Illumination and microscopy imaging

For initial proof-of-concept experiments, we used DLP patterned light to illuminate HeLa cells with high spatial resolution (Fig. 1c,d, Supplementary Fig. 2a,b, and Supplementary Video 1-2). To generate the spatially patterned photomask seen in Supplementary Figure 2a (left), a UCSB logo was saved as a binary bmp file and imported into Nikon Elements software (version 5.21.02). The photomask was then DMD projected over a HeLa monolayer labeled with an ‘inner’ Cy3 and ‘outer’ Cy5 conjugated PHO (Supplementary Fig. 1b – scheme (i)) for an 18-second time lapse with 1 second intervals (Supplementary Video 1). We used Nikon Elements software to generate the linear gradient seen in Supplementary Figure 2b (left), which followed the same illumination approach as above, for a 23-second time lapse (Supplementary Video 2). Complete photocleavage of PHO molecules was achieved around 25- 30 seconds using DLP illumination (Supplementary Fig. 2c). An LED floodlight was used in the proof-of-concept experiment (Fig. 1f,g and Supplementary Fig. 3a) and the integration of MPO experiment (Fig. 1h) to provide a convenient alternative for greater illumination area of cell samples. The HeLa cells in the proof-of-concept experiment (Fig. 1f,g and Supplementary Fig. 3), were labeled with an ‘inner’ Cy3 and ‘outer’ Cy5 conjugated PHO (Supplementary Fig1b – scheme (i)), imaged for Cy3 and Cy5 fluorescence, exposed to 0, 1, or 10 minutes of floodlight illumination, and then imaged again in each channel. Complete photocleavage of PHO molecules was achieved around 10 minutes using floodlight illumination (Supplementary Fig. 3b).

For the HeLa-mCherry and wildtype U2OS coculture experiment (Supplementary Fig. 4), cocultures were seeded in multiple wells of a glass-bottom 96-well plate. The cocultures were incubated with an ‘outer’ Cy5 conjugated PHO (Supplementary Fig1b – scheme (ii)), thus labeling both cells (Supplementary Fig. 4a). To specifically target HeLa-mCherry cells for light exposure, we employed a DLP light patterning strategy as follows. First, a comprehensive scanning of the entire area of multiple wells of a 96-well plate for mCherry fluorescence was performed and customized ROI photomasks were generated simultaneously for each tile by applying a threshold on mCherry fluorescence using Nikon Elements software. Next, HeLa cells were exposed to light delivered via DLP illumination for a duration of 30 seconds (Supplementary Fig. 4a). We implemented a 10% overlap for all x-y positions within the total area of each well to ensure comprehensive coverage. To ensure proper photocleavage of HeLa cells, and not U2OS cells, we imaged Cy5 fluorescence before and after illumination (Supplementary Fig. 4a).

For the MPO experiment (Fig. 1h), HeLa and U2OS cells were seeded in four separate wells of a glass-bottom 96-well plate. Each well was incubated with an ‘outer’ Cy5 conjugated PHO (Supplementary Fig. 1b – scheme (ii)) as well as a unique MPO. Both cell types were then imaged for Cy5 fluorescence, illuminated with no light or 1 minute of light, imaged again, and then processed for sequencing.

For the error correction experiment (Fig. 1i-k), HeLa cells were seeded in separate wells of a glass-bottom 96-well plate for each light treatment. Each well was incubated with one unique ‘outer’ Cy5 conjugated PHO (Supplementary Fig1b – scheme (ii)) and five unique non fluorophore conjugated PHOs (Supplementary Fib1b – scheme (iii)). We employed DLP illumination for various specified durations of light exposure where the entire area of each well was given a different light dose (Fig. 1i and Supplementary Fig. 8a,b). We implemented a 10% overlap for all x-y positions within the total area of each well to ensure comprehensive coverage. To ensure proper photocleavage, we imaged Cy5 fluorescence before and after illumination.

For the combinatorial barcoding experiment (Fig. 2a-d and Supplementary Fig. 9), HeLa cells were seeded in multiple wells of a circular 1000 µm diameter fibronectin-coated protein-patterned 96-well plate (CYTOO, 20-950-00) to maintain all cells within the field of view of a 10x objective. The cell monolayer was then subjected to three rounds of: incubation with a uniquely fluorophore conjugated and barcoded PHO (Supplementary Fig1b – scheme (ii)), imaging for the fluorescence channels corresponding to PHO1-3, illumination of a binary photomask for 30 seconds, wash to remove cleaved and unbound PHO molecules, and repeated imaging for the fluorescence channels corresponding to PHO1-3 post illumination (Supplementary Fig. 9a).

For the experiments determining the adaptability of scSTAMP-seq with droplet-based single cell sequencing (Fig. 2e,f and Supplementary Fig. 11a,b), and simultaneous multiomics sequencing with scSTAMP-MAT-seq (Fig. 2g,h, Supplementary Fig. 13,14), we used floodlight illumination, as patterned light was not necessary for determining the feasibility of the aspects above, and allowed for a larger sample of cells to be illuminated in a more convenient time frame than can be achieved via DMD light projection. For the droplet-based single cell sequencing experiment (Fig. 2e,f and Supplementary Fig. 11 a,b), HeLa cells were seeded in multiple wells of a 96-well plate, incubated with an ‘outer’-Cy5 conjugated PHO (Supplementary Fig1b – scheme (ii)), imaged for Cy5 fluorescence, exposed to 0, 1, or 10 minutes of floodlight illumination, and imaged again for Cy5 fluorescence post illumination. For the scSTAMP-MAT-seq experiment (Fig. 2g,h, Supplementary Fig. 13,14), HeLa and U2OS cells were seeded in four separate wells of a glass-bottom 96-well plate. Each well was incubated with an ‘outer’ Cy5 conjugated PHO (Supplementary Fig1b – scheme (ii)), imaged for Cy5 fluorescence, illuminated with no light or 1 minute of light, imaged again, and then processed for sequencing.

### Image processing and analysis

All images were generated as Nikon nd2 files, processed on ImageJ/FIJI, converted to tiff files, and analyzed using custom code on MATLAB (version R2022a) and Python (version 3.9.0). Representative figures were generated in MATLAB, R (version 4.2.2), and RStudio (version 2022.12.0+353). Representative images and figures were compiled in Adobe Illustrator (version 26.5).

For initial experiments that employed spatial DMD light patterning (Fig. 1c,d), composite multi-channel fluorescence images of Cy3 and Cy5 fluorescence were captured over three z-planes spaced in 15 µm intervals. The z slices for each individual channel were blended using an applied maximum intensity projection (maxIP) on Nikon Elements software. Each channel was separated using the ‘split channels’ function in ImageJ/FIJI. Brightness and contrast LUTs (lookup tables) for the minimum and maximum intensity were manually selected for the first frame of each channel to enhance visualization. The adjusted LUTs were then applied to each frame of the time series. Image 8 of the 18-image and 23-image time series for grayscale Cy3 and Cy5 fluorescence after applying DMD light patterns are shown in Supplementary Fig. 2a,b, respectively. ‘Blue’ and ‘red hot’ false color LUTs were then applied for Cy3 and Cy5, respectively, to enhance color contrast between the two fluorescence channels. The two channels were then merged using the ‘merge channels’ function in ImageJ/FIJI to generate the images for Figure 1c,d and the videos for Supplementary Videos 1-2.

The Cy3 and Cy5 fluorescence within the gradient ROI were analyzed across the 23- image time stack using custom code written on MATLAB. A maximum threshold and background subtraction was applied to each image to remove saturated pixels. Fluorescence intensity was then normalized for each stack of the 2 channels. The change in intensity between Cy3 and Cy5 within the gradient was calculated over the 23 second time course by taking Cy5 fluorescence over Cy3 fluorescence.

Experiments after the proof-of-concept experiments (Fig. 1c,d) were processed similarly as above. However, for future experiments, only 1 z-plane (opposed to 3) was captured to reduce imaging time, so the maximum intensity projection processing step was omitted. Additionally, in future iterations of experiments, we opted for ‘outer’ fluorophore conjugated PHOs (opposed to ‘inner’ fluorophore and ‘outer’ fluorophore conjugated PHOs) and thus captured and analyzed one channel for PHO fluorescence.

For images in the coculture experiment (Supplementary Fig. 4a), single cells were segmented by implementing the Cellpose2.0 ‘cyto2’ model on ‘outer’ Cy5 conjugated PHO fluorescence before photocleavage using a custom code on Python. The mean fluorescence value for each segmented single cell was calculated for 69 frames in the 96-well (Supplementary Fig. 4b).

For the combinatorial barcoding experiment (Supplementary Fig. 9), a 75%, 50%, and 25% area photomask were applied at each subsequent round (Supplementary Fig. 9a top row). Round 1 contained an ‘outer’ Cy5 conjugated PHO1 (Supplementary Fig. 9a – middle left), round contained an ‘outer’ FAM conjugated PHO2 (Supplementary Fig. 9 – middle center), and round contained an ‘outer’ Cy3 fluorophore conjugated PHO3 (Supplementary Fig. 9a – middle right). Fluorescence channels were false colored magenta, yellow, and cyan, for PHOs 1, 2, and 3, respectively, and then overlayed to visualize the combinations of PHO fluorescence after each step (Supplementary Fig. 9a bottom row). To analyze images for the combinatorial barcoding experiment, single cells were manually segmented by drawing ROIs around individual cells in ImageJ/FIJI. 40 cells were selected within each quadrant for 120 cells total. The median fluorescence value for the segmented single cells in each quadrant of a protein-patterned monolayer was calculated for the three PHO fluorescence channels using custom code on MATLAB (Supplementary Fig. 9b).

### Sample collection and single-cell processing

Single-cell suspensions were made by dissociating cells using 0.25% trypsin-EDTA and inactivated using DMEM containing serum. Afterwards, cells were washed and resuspended in 1 mL of cold 1x DPBS-/- (without calcium or magnesium) (Gibco, 14040133) before being passed through a 35 µm nylon mesh cell strainer into a 5 mL round bottom polystyrene tube (Corning, 352235) that was then kept on ice. For plate-based single-cell capture, cell suspensions were FACS sorted for single cells into 384-well plates (Bio-Rad, HSP3801). For droplet-based single-cell capture, cell suspensions were processed for single cells by a 10X Genomics Chromium Controller following the workflow of the 10X Genomics Chromium Next GEM Single Cell 3’ LT Kit v3.1 (10X Genomics, 1000325).

### scSTAMP-seq library preparation

*scSTAMP-seq plate-based library preparation*. 4 µL of Vapor-Lock (QIAGEN, 981611) was dispensed into each well of a 384-well plate using a 12-channel pipette. All downstream dispensing into 384-well plates was performed using a Nanodrop II liquid handling robot (BioNex Solutions). To each well, 100 nL of uniquely barcoded 7.5 ng/µL reverse transcription primers containing a 6-nucleotide UMI was added. The reverse transcription primers used here have been described by Grün *et al.*^33^. Next, 100 nL of lysis buffer (0.175% IGEPAL CA-630, 1.75 mM dNTPs, 1:1,250,000 ERCC RNA spike-in mix (Ambion, 4456740), 0.19 U RNase inhibitor (Clontech, 2313A)) was added to each well. Single cells were sorted into individual wells of a 384-well plate using FACS. After sorting, plates were heated to 65°C for 3 minutes and returned to ice.

Reverse transcription was conducted next by adding 150 nL of reverse transcription mix (0.7 U RNase OUT (Invitrogen, 10777-019), 2.34x first strand buffer (Invitrogen, 10777-019), 23.34 mM DTT (Invitrogen, 10777-019), 3.5 U Superscript II (Invitrogen, 18064-071)) to each well and the plates were incubated at 42°C for 1 hour and 15 minutes, 4°C for 5 minutes, 70°C for 10 minutes, and then held at 4°C. Next, second strand synthesis was carried out by adding 1.75 µL of second strand synthesis mix (1.74x second strand buffer (Invitrogen, 10812-014), 0.35 mM dNTP (Invitrogen, 10812-014), 0.14 U *E. coli* DNA Ligase (Invitrogen, 18052-019), 0.56 U *E. coli* DNA Polymerase I (Invitrogen, 18010-025), 0.03 U RNase H (Invitrogen 18021-071)) to each well and the plates were incubated at 16°C for 2 hours and then held at 4°C. 1 µL of nuclease-free water was added into each well to minimize losses during downstream pooling. The PHO, MPO, and endogenous mRNA molecules in a well now all contain the same RNA capture barcode, indicating a unique cell, which allows for pooling. Reaction wells receiving different barcodes were pooled using a multichannel pipette, and the oil phase was discarded.

The small molecule spatial barcode fraction was separated from endogenous transcript cDNA using a 3.2x solid phase reversible immobilization (SPRI) size selection with 1.8x 100% isopropanol in 2 steps. First, 0.7x AMPure XP paramagnetic SPRI beads (Beckman Coulter, A63881) were added and incubated at room temp for 15 minutes. At this step, it is critical to not dispose of the supernatant, as the spatial barcode library fraction is found in the supernatant and the endogenous cDNA is bound to SPRI beads. The samples were placed on a magnetic stand for 5 minutes and the supernatant containing the spatial barcode fragments was then transferred to a fresh Eppendorf tube. The endogenous cDNA fraction was prepared as described previously by Chialastri *et al.*^31^. Second, to the spatial barcode sample, an additional 2.5x SPRI beads and 1.8x 100% isopropanol (of the original pre-cleanup sample volume) were added and incubated at room temp for 15 minutes. The samples were placed on a magnetic stand for 5 minutes. The supernatant was removed and beads were washed twice with 80% ethanol. The sample was then eluted in 30 µL of nuclease-free water. The sample was then concentrated to 15 µL using an Eppendorf Vacufuge plus. Next, samples were prepared for PCR amplification by mixing 15 µL of the concentrated elute from the previous step, 25 µL of 2x PCR master mix (NEB, M0541S), 2 µL Illumina RPI primer (10 µM), 2 µL each of an ‘inner’ and ‘outer’ PCR primer (10 µM), and nuclease-free water up to 50 µL. This first PCR was performed at 94°C for 10 seconds, 60°C for 30 seconds, and 72°C for 30 seconds, for 10 cycles. A subsequent ExoSAP-IT treatment was then performed to enrich the PCR product as follows. 2 µL of ExoSAP-IT (Applied Biosystems, 78200.200.UL) was added to 5 µL of sample from the previous PCR (for a total volume of 7 µL) and incubated at 37°C for 15 minutes. Another PCR was then performed by mixing 7 µL of the enriched PCR sample with 22.5 µL 2x PCR master mix, 2 µL Illumina RPI primer (10 µM), 2 µL Illumina sample index primer (10 µM), and nuclease-free water up to 45 µL. This second PCR was performed at 94°C for 10 seconds, 60°C for 30 seconds, and 72°C for 30 seconds, for 6-12 cycles. Another ExoSAP-IT treatment was then performed to enrich the 45 µL reaction by adding 5 µL of ExoSAP-IT to the previous PCR (for 50 µL total) and incubated at 37°C for 15 minutes. Finally, a 1.2x double-sided selection (0.90x left/0.30x right) was used to purify the spatial library, and a final 80% ethanol wash was performed twice. The product was then eluted in 20 µL of nuclease-free water. Library fragment size distribution was determined using a 2100 Bioanalyzer. A schematic depicting each step of this process can be found in Supplementary Figure 1c.

#### scSTAMP-seq droplet-based library preparation

Sequencing libraries were prepared using a custom protocol based on the 10X Genomics Chromium Next GEM Single Cell 3’ LT Kit v3.1 (10X Genomics, 1000325) adapted from McGinnis *et al.*^29^. The 10X workflow was followed until cDNA amplification. We designed custom ‘inner’ and ‘outer’ PHO handle primers that allow scSTAMP- seq to be adapted for the 10X workflow (Supplementary Table 1). The scSTAMP-seq 10X ‘inner’ and ‘outer’ PHO primers were mixed to a working concentration of 2.5 µM. Next, the following cDNA amplification master-mix was made: 50 µL 10X amplification master mix, 15 µL 10X cDNA primer mix, 1 µL scSTAMP-seq 10X primer mix. Then, cDNA amplification was performed and an SPRI size selection was performed by incubating samples for 30 minutes with 0.6x AMPure XP paramagnetic SPRI beads to separate spatial and endogenous cDNA fractions. Next, the small molecule spatial barcode fraction was separated from endogenous transcript cDNA using the same strategy as described in the plate-based library preparation method above. The endogenous cDNA fraction was then processed to generate Illumina libraries according to the 10X workflow and the spatial fraction was processed to generate Illumina libraries as described in the plate-based library preparation workflow. A schematic depicting each step of this process can be found in Supplementary Figure 10.

#### scSTAMP-MAT-seq multiomic library preparation

The DNA accessibility and DNA methylation sequencing pipeline was performed as described previously in the scMAT-seq methods by Chialastri *et al.*^31^ up to the first SPRI bead clean up. At this step, the small molecule spatial barcode fraction was separated from endogenous molecules using the same strategy as described in the plate-based library preparation workflow. The transcriptomic and epigenomic libraries were processed to generate Illumina libraries as described in scMAT-seq by Chialastri *et al.*^31^ and the spatial fraction was processed to generate Illumina libraries as described in the plate-based library preparation workflow above. A schematic depicting each step of this process can be found in Supplementary Fig. 12.

Library sequencing was performed by Novogene on an Illumina HiSeq 4000 platform or at the Biological Nanostructures Laboratory (BNL) in the California NanoSystems Institute (CNSI) at the University of California Santa Barbara (UCSB) on an Illumina NextSeq 500 platform.

### scSTAMP-seq data processing and gene expression analysis

Sequencing FASTQ data files were processed using custom Perl scripts on a Linux high performance computing (HPC) cluster at the University of California, Santa Barbara. Sequencing data was post-processed and analyzed using Seurat (version 4.3.0), R (version 4.2.2), and RStudio (version 2022.12.0+353). Representative figures were generated in RStudio and compiled in Adobe Illustrator (version 26.5).

#### scSTAMP-seq barcode analysis

A custom Perl script was written to evaluate the spatial barcode fraction of the sequencing library. For PHO spatial barcode sequences, read 1 was used to determine cell barcode/UMI. Read 2 was used to distinguish ‘inner’ and ‘outer’ sequence reads by identifying unique ‘inner’ and ‘outer’ primer sequences as well as the downstream spatial barcode/UMI sequence. MPO barcode counts were determined similarly, but an additional multiplex identifier sequence from read 2 was used to distinguish multiplex from spatial barcodes. Unique counts for each molecule were defined as reads containing a unique combination of the UMI on read 1 and on read 2.

#### Noise reduction through PHO multiplexing

For the error correction experiment (Fig. 1i-k and Supplementary Fig. 8), the following steps were used to correctly assign light dosage to cells with ambiguous ratios of PHO cleavage: (1) For each illumination level, the distribution of ‘outer’-to- ‘inner’ PHO cleavage ratios was used to define an upper and lower threshold at 10% and 90%, and the region of the distribution within these thresholds is referred to as the effective region. Cells with PHO cleavage ratios within the effective region of multiple adjacent illumination levels were labeled as ambiguous. Regions not assigned in any effective region were corrected by the nearest two effective regions by equally extending them to cover the unassigned region. (2) For each cell, the ‘outer’-to-‘inner’ cleavage ratio of each unique PHO was compared to the effective region for all illumination levels to identify which illumination level a cell corresponds to. A cell can be assigned to multiple illumination levels as there exists overlap between the effective region of different illumination levels. This step was repeated for all unique PHOs. (3) By combining data from unique PHOs on a cell, the counts of illumination levels that correspond to a cell were calculated. For each cell, the illumination level with the highest counts was assigned as the predicted illumination level that a particular cell was exposed to. The predicted result was then compared to the actual illumination level to calculate the percent accuracy in spatial assignment. If more than one illumination level had the highest count for a cell, it was considered an incorrect prediction.

#### scSTAMP-seq gene expression analysis

For plate-based sequencing, read 2 of the paired-end read was aligned in the sense direction using BWA (version 0.7.15-r1140) to the RefSeq gene model based on the human genome release hg19, incorporating a collection of 92 ERCC spike-in molecules. In cases where a read aligned to multiple locations, it was evenly distributed across those locations. Gene isoforms were consolidated into a single gene count, and the unique molecular identifiers (UMIs) were used to remove duplicate reads and generate transcript counts at the single-molecule level for each gene within individual cells. Genes that were not detected in at least one cell were excluded from subsequent analyses. For droplet-based sequencing, read 2 was trimmed using the default settings of TrimGalore. After trimming, STARsolo (STAR aligner version 2.7.8a) was used to map the reads to hg19 using the gene annotation file from Ensembl. Transcripts with the same UMI were deduplicated and genes that were not detected in at least one cell were removed from any downstream analysis.

The standard analysis pipeline in Seurat (version 4.3.0) was used for single-cell RNA expression normalization and analysis^36^. For plate-based sequencing experiments, cells containing more than 1000 genes and more than 1000 unique transcripts, as well as less than 20% ERCC spike-ins, were used for downstream analysis. For the droplet-based sequencing experiment, cells containing more than 200 genes and more than 200 unique transcripts, as well as less than 5% mitochondrial counts, were used for downstream analysis. The default NormalizeData function was used to log normalize the data. Principal components were obtained from the 2,000 most variable genes and the elbow method was used to determine the optimal number of principal components used in clustering. UMAP based clustering was performed by running the following functions, FindNeighbors, FindClusters, and RunUMAP. After clustering, cell types were assigned to groups using known expression markers, spatial barcode counts, and multiplex barcode counts, where stated.

### scSTAMP-MAT-seq genome-wide DNA accessibility and DNA methylation analysis

The scSTAMP-MAT-seq analysis pipeline for DNA accessibility and 5mC was performed as described in the scMAT-seq methods by Chialastri *et al.*^31^. DNase I hypersensitivity sites for HeLa cells were downloaded from the UCSC table browser and sites were grouped based on detection scores^35^. To compare across datasets, the data was normalized to counts per million and a 75 base pair moving average was plotted for region of interest. When comparing differing genomic regions within a sample, each region was further normalized by methylated cytosines that were detected when MspJI-seq was performed on bulk HeLa gDNA that had been GpC methylated after stripping off chromatin. Bulk HeLa gDNA isolation, chromatin stripping, and processing was performed as described previously by Chialastri *et al.*^31^.

For clustering, DNA accessibility and DNA methylation were quantified within 5 kb bins and then converted to binary scores. Pseudobulk profiles were generated using the assigned cell type from the transcriptome. For comparison between HeLa and U2OS cells, the top 2% most variable bins between groups were retained. After removal of bins with low variance, principal component analysis was used to assign clusters. Cluster identification was performed through comparison to transcriptome derived cell types, where high similarity was observed between cluster calling for all 3 measurements. A threshold minimum of 15,000 was applied to the data shown in Supplementary Figure 14a,b.

## Acknowledgements

We thank members of the Dey and Wilson labs for helpful discussions. We thank Jennifer Smith at the Biological Nanostructures Laboratory in the California NanoSystems Institute (CNSI), supported by UCSB and UC Office of the President, for help with Illumina sequencing, and Travis Parsons at Nikon Instruments for assistance in microscopy. We thank the Acosta-Alvear lab for providing HeLa-mCherry cells. Computational work was supported by the Center for Scientific Computing at CNSI and Material Research Laboratory (MRL) at UCSB: an NSF MRSEC (DMR-1720256) and NSF CNS-1725797. This work was supported by the Arnold and Mabel Beckman Foundation Scholars Award to M.G., the CIRM Research Training Grant EDUC4-12821 to R.A.P., the U.S. Army Research Office contract W911NF-19-D-0001 and cooperative agreement W911NF-19-2-0026 for the Institute for Collaborative Biotechnologies to M.Z.W., and the NIH grants R01HD099517, R01HG011013 and an NSF grant #2339849 to S.S.D.

## Author contributions

Conceptualization, R.A.P., A.C., M.Z.W, S.S.D.; Methodology, R.A.P., A.C., M.Z.W., S.S.D.; Investigation, R.A.P., A.C., C.W., M.G., W.W.; Formal Analysis, R.A.P., A.C., C.W., M.G., K.H.; Writing – Original Draft, R.A.P.; Writing – Review & Editing, R.A.P., A.C., C.W., M.G., M.Z.W., S.S.D.; Funding Acquisition, M.Z.W., S.S.D.; Resources, M.Z.W., S.S.D.; Supervision, M.Z.W., S.S.D.

## Data availability

Sequencing data have been deposited in the Gene Expression Omnibus (GEO) database accession code GEO: GSE237524.

## Code availability

Custom code for analyzing scSTAMP-seq data and the accompanying documentation are available on GitHub at https://github.com/deylabucsb/scSTAMP-seq.

