## Supplementary Information for "Photolabile oligonucleotides combined with topologically imposed light gradients enable spatially resolved single-cell transcriptomics and epigenomics"

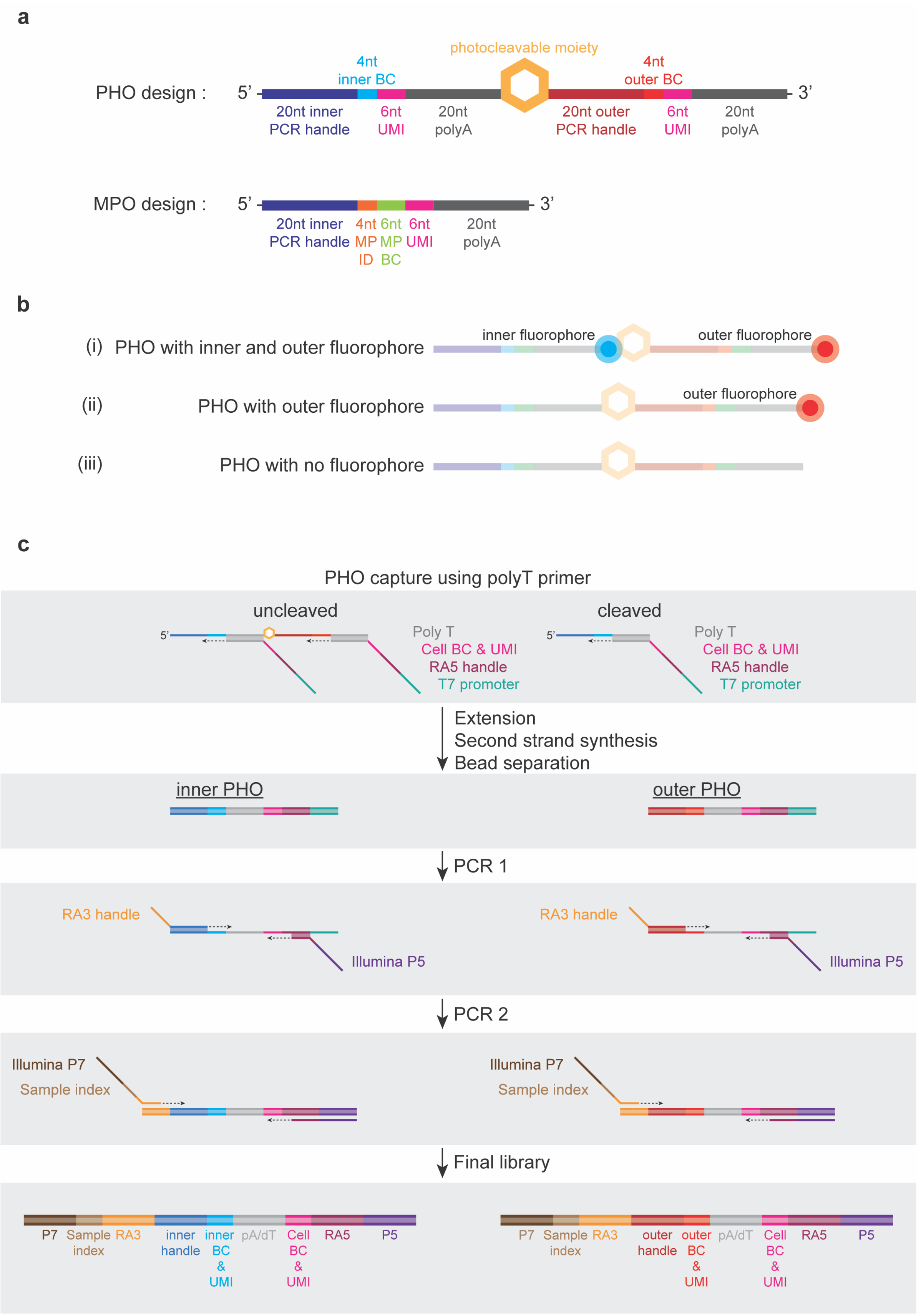
a
PHO design :
5'
3'
4nt
inner BC
photocleavable moiety
4nt
outer BC
20nt inner
PCR handle
6nt
UMI
20nt
polyA
20nt outer
PCR handle
6nt
UMI
20nt
polyA
MPO design :
5'
3'
20nt inner
PCR handle
4nt
MP
ID
6nt
MP
BC
6nt
UMI
20nt
polyA
b
(i) PHO with inner and outer fluorophore
inner fluorophore
outer fluorophore
(ii) PHO with outer fluorophore
outer fluorophore
(iii) PHO with no fluorophore
c
PHO capture using polyT primer
uncleaved
cleaved
Poly T
Cell BC & UMI
RA5 handle
T7 promoter
Poly T
Cell BC & UMI
RA5 handle
T7 promoter
Extension
Second strand synthesis
Bead separation
inner PHO
outer PHO
PCR 1
RA3 handle
Illumina P5
RA3 handle
Illumina P5
PCR 2
Illumina P7
Sample index
Illumina P7
Sample index
Final library
P7
Sample index
RA3
inner handle
inner BC & UMI
pA/dT
Cell BC & UMI
RA5
P5
P7
Sample index
RA3
outer handle
outer BC & UMI
pA/dT
Cell BC & UMI
RA5
P5

**Supplementary Figure 1 | Design of different PHO variants and downstream capture in single-cell sequencing.** **a**, Schematic depicts the components that comprise the 'inner' and 'outer' elements of PHOs and MPOs. Both 'inner' and 'outer' DNA sequences of PHOs contain a handle for downstream PCR amplification, an 'inner'- and 'outer'-specific barcode, a 6-nucleotide UMI to count individual molecules, and a poly-A capture sequence. The 'inner' PCR handle is also used to anneal to a complementary sequence on the CMO anchor for embedding in cell membranes. MPOs resemble the 'inner' PHO sequence but contain an additional multiplex identification sequence (MP ID) to distinguish MPOs from PHOs. **b**, Schematic shows three PHO fluorophore conjugation schemes. PHOs can be conjugated with distinct fluorophores to enable visualization of cleavage of the 'outer' sequence prior to downstream sequencing. **c**, Schematic of PHO library preparation. First, uncleaved and cleaved PHOs are copied using a primer that contains a T7 promoter (teal), an RA5 handle (purple), a cell specific barcode, a UMI (magenta), and a polyT capture sequence (gray). Second strand synthesis is subsequently performed, and PHO derived molecules are separated from endogenous cDNA using SPRI beads. Next, an RA3 handle primer (yellow) and an 'inner' (blue) or 'outer' (red) adapter, along with an RA5 adapter with an Illumina P5 (indigo) primer are added to amplify and extend PHO molecules. Finally, a primer containing an Illumina P7 (brown), a sample index sequence (tan) and an RA3 adapter, along with the same RA5-P5 adapter primer in the previous step, are used to amplify and extend PHO molecules to generate final Illumina libraries.

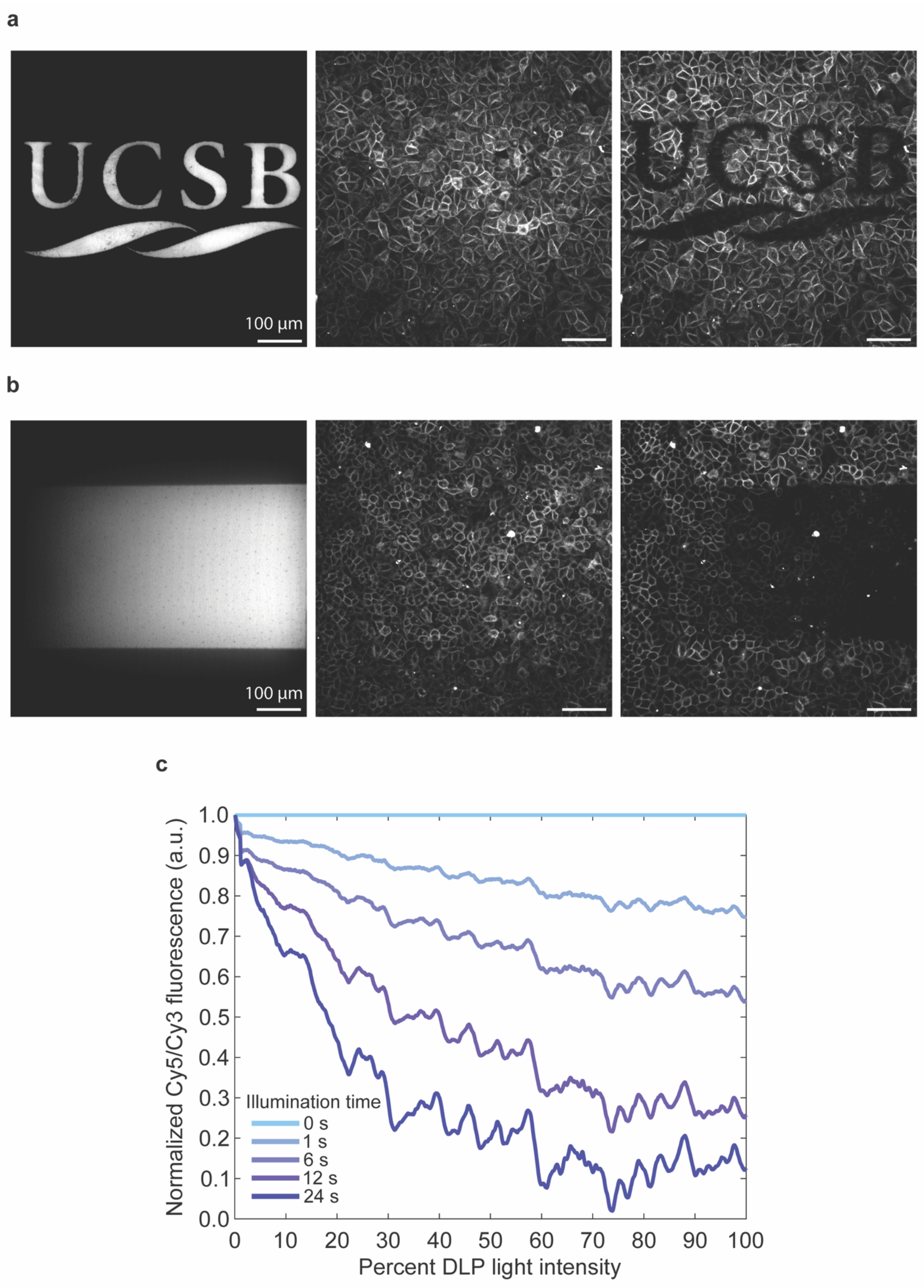

a
UCSB
100 µm
b
100 µm
c
Normalized Cy5/Cy3 fluorescence (a.u.)
1.0
0.9
0.8
0.7
0.6
0.5
0.4
0.3
0.2
0.1
0.0
0
10
20
30
40
50
60
70
80
90
100
Percent DLP light intensity
Illumination time
0 s
1 s
6 s
12 s
24 s

**Supplementary Figure 2 | User defined DMD illumination over live cells. a**,**b**, Images of DMD-projected photomasks (left) for a binary UCSB logo (a) and a linear gradient (b) used to illuminate live cells labeled with an 'inner' Cy3 and 'outer' Cy5 conjugated PHO. Images of 'inner' Cy3 (middle) and 'outer' Cy5 (right) fluorescence post illumination. **c**, Normalized fraction of Cy5 over Cy3 fluorescence as a function of percent DLP light intensity of the linear gradient for specified exposure times in the time lapse shown in Supplementary Video 2.

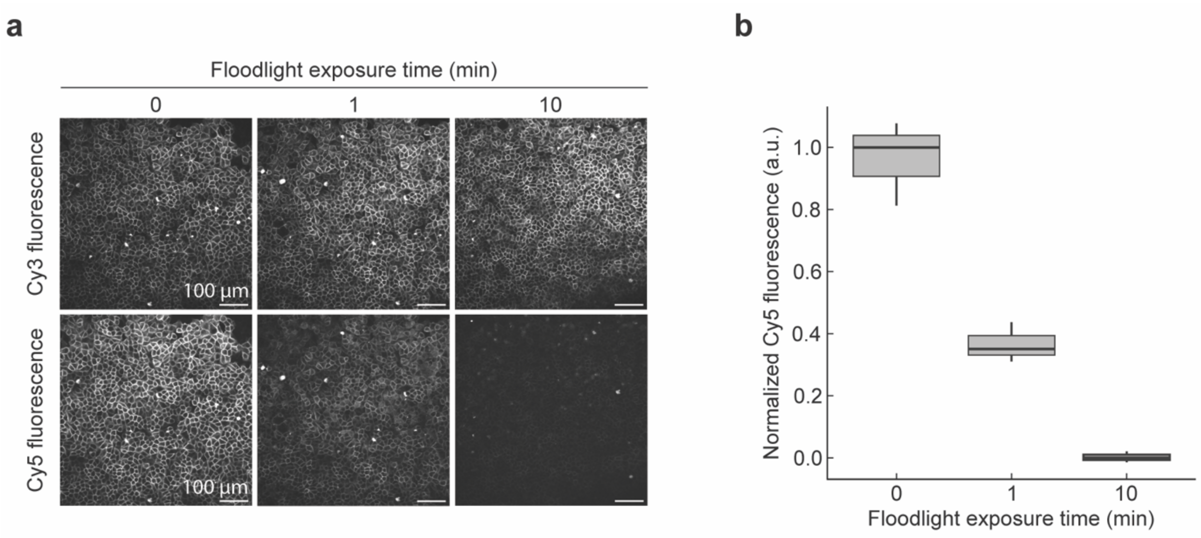


**Supplementary Figure 3 | LED floodlight illumination over live cells. a**, Fluorescence images of live HeLa cells labeled with PHOs containing an 'inner' Cy3 fluorophore (top) and 'outer' Cy5 fluorophore (bottom) for three distinct durations of illumination with a 365-nm LED floodlight. **b**, Boxplot of normalized Cy5 fluorescence shown in (a) corresponding to the three different light exposure times.

**a**

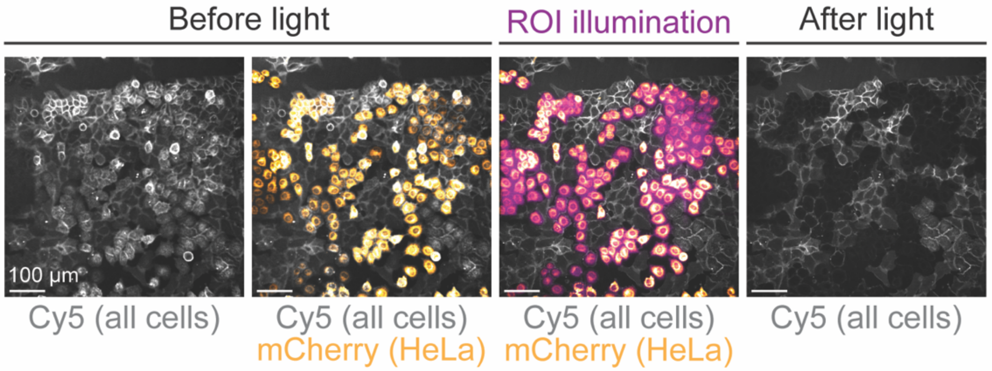


**b**

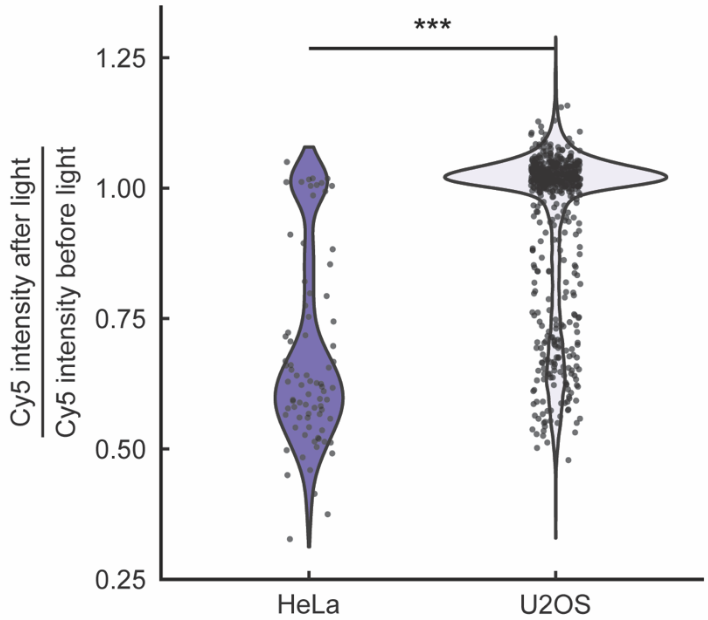


**Supplementary Figure 4 | scSTAMP-seq can be used to label individual cells. a**, Composite fluorescence images of an 'outer' Cy5 conjugated PHO (gray) labeled coculture containing wildtype U2OS and mCherry expressing HeLa cells (orange). Illuminated photomasks (magenta) were generated by thresholding on mCherry positive HeLa cells. **b**, Violin plot of the ratio of Cy5 fluorescence after and before light exposure for HeLa and U2OS cells. *** indicates a statistically significant difference between distributions ($p$ < 2.22e-16, Mann-Whitney U test).

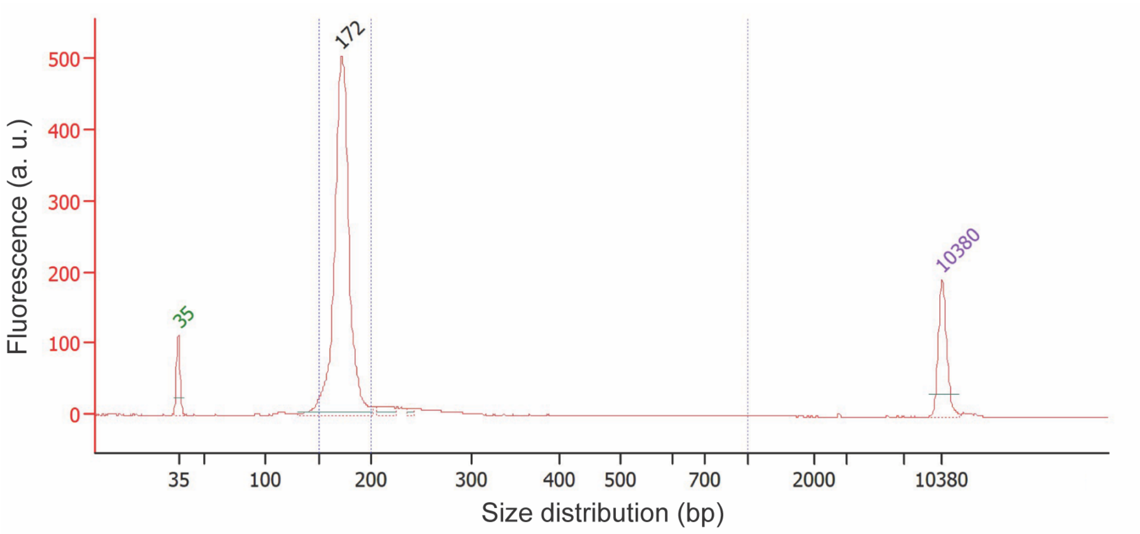


**Supplementary Figure 5 | Bioanalyzer trace of spatial PHO Illumina library in scSTAMP-seq.** Bioanalyzer traces shows the expected size of the PHO Illumina library after enrichment with 3.2x SPRI beads with 1.8x 100% isopropanol following the amplification protocol described in Supplementary Figure 1c.

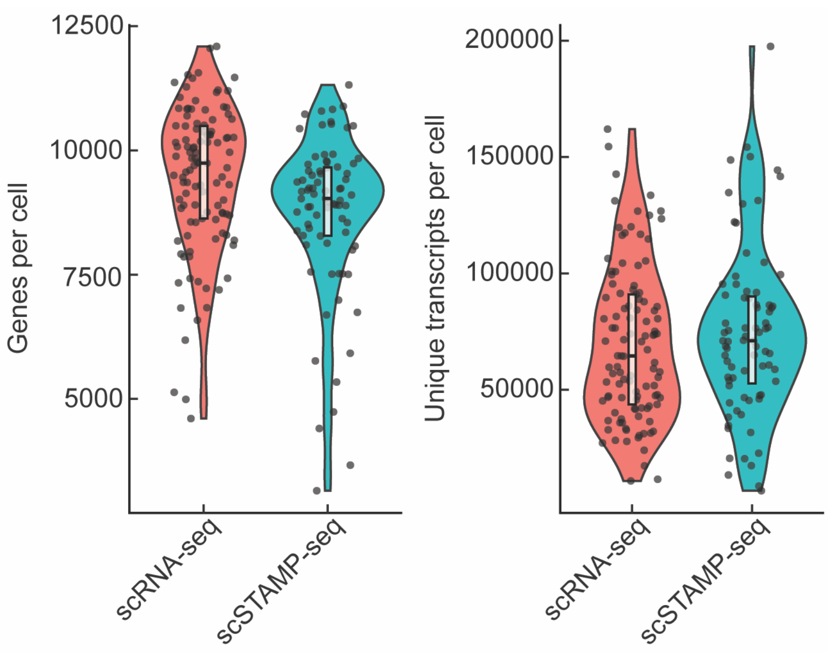


**Supplementary Figure 6 | Comparison of the transcriptome obtained from scSTAMP-seq to scRNA-seq.** Violin plots of the number of genes and total number of unique transcripts detected per cell for standard scRNA-seq and scSTAMP-seq.

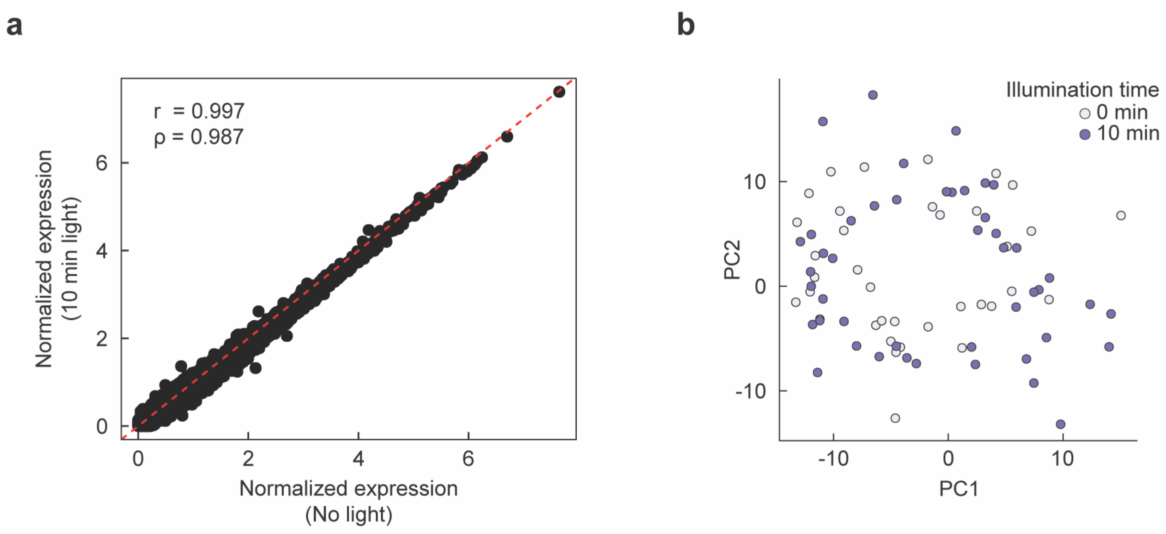


**Supplementary Figure 7 | UV illumination does not impact the endogenous transcriptome for plate-based cell capture and sequencing in scSTAMP-seq. a**, Scatterplot of the normalized gene expression levels of cells exposed to no light compared to saturating UV light required for full PHO cleavage (Pearson's $r$ = 0.997 , Spearman's $\rho$ = 0.987). **b**, Principal component projection of cells that received saturating light and no light treatment in scSTAMP-seq.

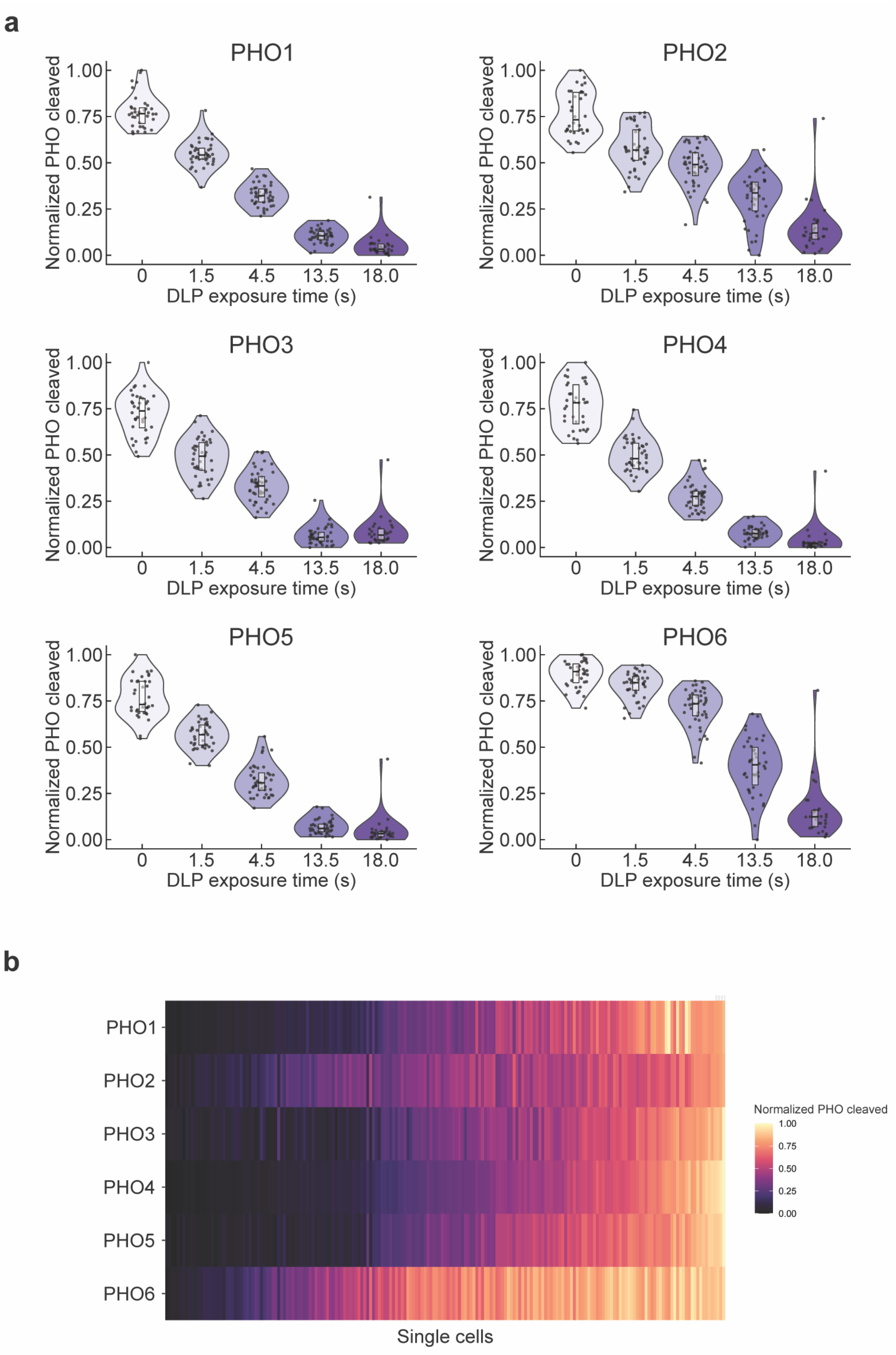


**Supplementary Figure 8 | PHO multiplexing enables error correction and improves the spatial resolution of scSTAMP-seq. a**, Violin plots of normalized ‘outer’-to-‘inner’ PHO sequencing read counts for PHO1-6 in individual cells after 5 distinct illumination exposure times.

PHO1 is also shown in Fig. 1i. **b**, Heat map of normalized ‘outer’-to-‘inner’ PHO sequencing read counts for PHO1-6 in individual cells.

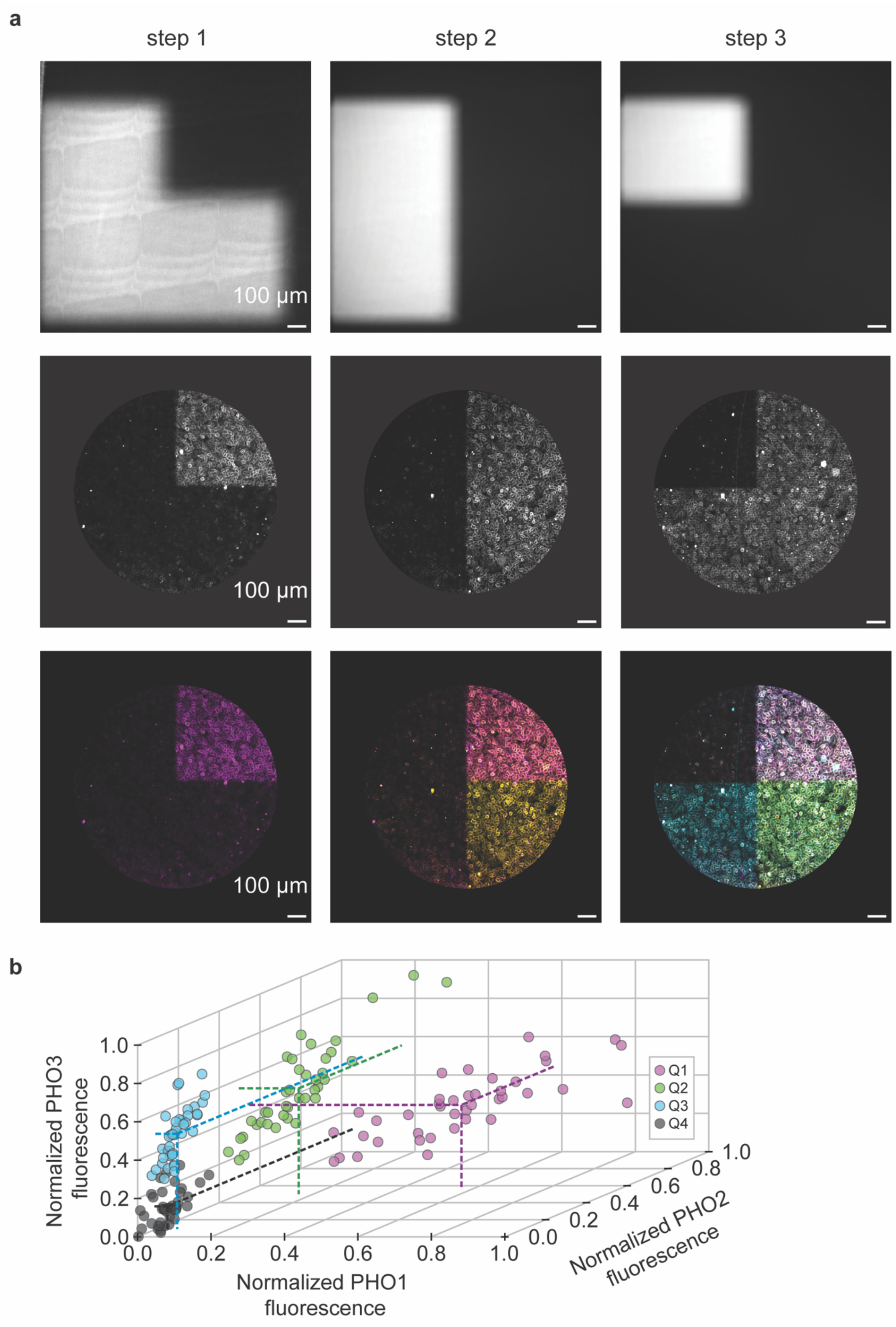
a
step 1
step 2
step 3
100 µm
100 µm
100 µm
b
Normalized PHO3 fluorescence
Normalized PHO1 fluorescence
Normalized PHO2 fluorescence
Q1
Q2
Q3
Q4
0.0
0.2
0.4
0.6
0.8
1.0

**Supplementary Figure 9 | Sequential rounds of optical labeling increases the spatial resolution of scSTAMP-seq. a**, Top row – images of the three binary photomasks applied sequentially to cells during the three rounds of optical labeling. Middle row – fluorescence images of cells labeled with Cy5 conjugated PHO1 (left), FAM conjugated PHO2 (middle), and Cy3 conjugated PHO3 (right) after photomask exposure. Bottom row – composite false-color overlap images of each PHO fluorophore at the three corresponding steps. **b**, 3D plot of the fluorescence intensity values of PHO1-3 for the image shown in Figure 2c. Individual cells are colored corresponding to assigned quadrant based on the combination of intensity values for each fluorophore. Dotted lines correspond to the average fluorescence for each unique PHO in each quadrant.

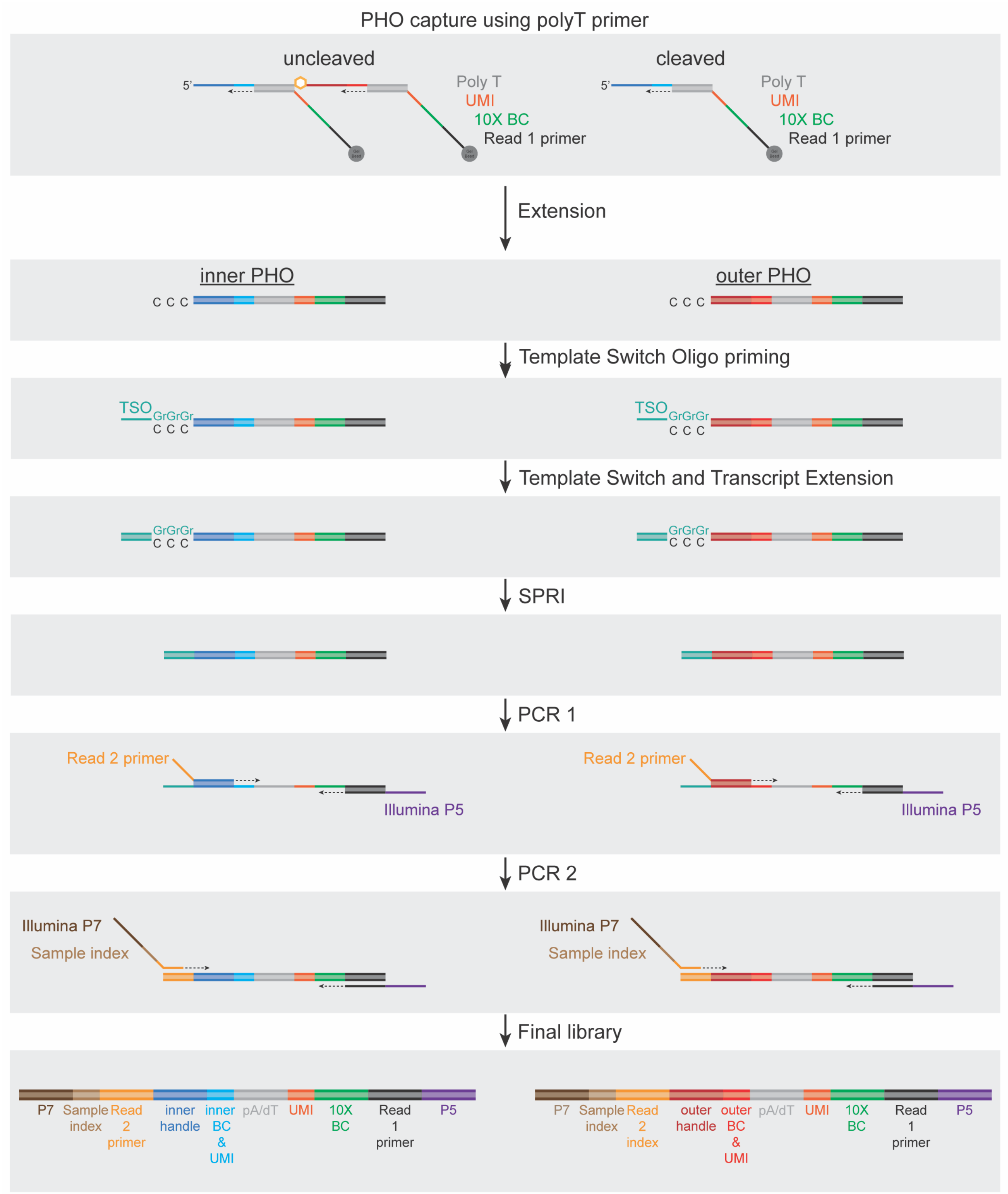


**Supplementary Figure 10 | Droplet-based sequencing library preparation in scSTAMP-seq.** Schematic of PHO library preparation on a droplet-based 10X platform. First, uncleaved and cleaved PHOs are copied using a polyT capture primer (gray) that also contains an Illumina TruSeq read 1 primer (black), a cell specific 10X barcode (green), and a UMI (orange). Next, a template switching oligo (TSO) (teal) is added and the molecules are extended. Prior to cDNA amplification, PHO derived molecules are separated from endogenous mRNA derived molecules using SPRI beads. Next, a TruSeq read 2 primer (yellow) with an 'inner' (blue) or 'outer' (red) adapter, along with an Illumina P5 (purple) with a read 1 primer, are added to amplify and extend PHO molecules. Finally, a primer containing an Illumina P7 (brown), a sample index sequence (tan) and a read 2 adapter, along with the same read 1-P5 adapter primer in the previous step, are used to amplify and extend PHO molecules to generate final Illumina libraries.

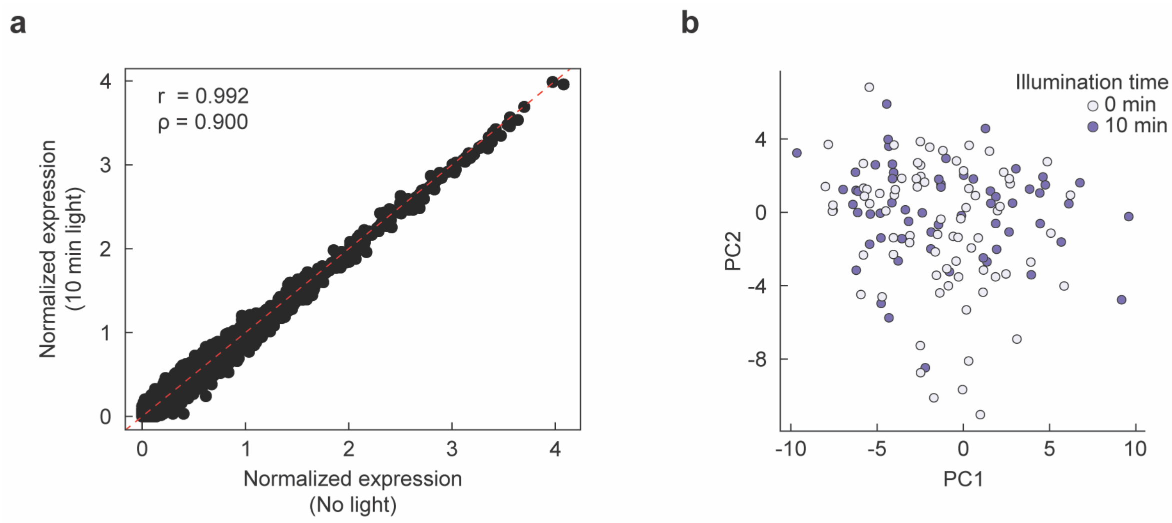


**Supplementary Figure 11 | UV illumination does not impact the endogenous transcriptome for droplet-based cell capture and sequencing in scSTAMP-seq.** **a**, Scatterplot of the normalized gene expression levels of cells exposed to no light compared to saturating UV light required for full PHO cleavage (Pearson's $r$ = 0.992, Spearman's $\rho$ = 0.900). **b**, Principal component projection of cells that received saturating light and no light treatment in scSTAMP-seq.

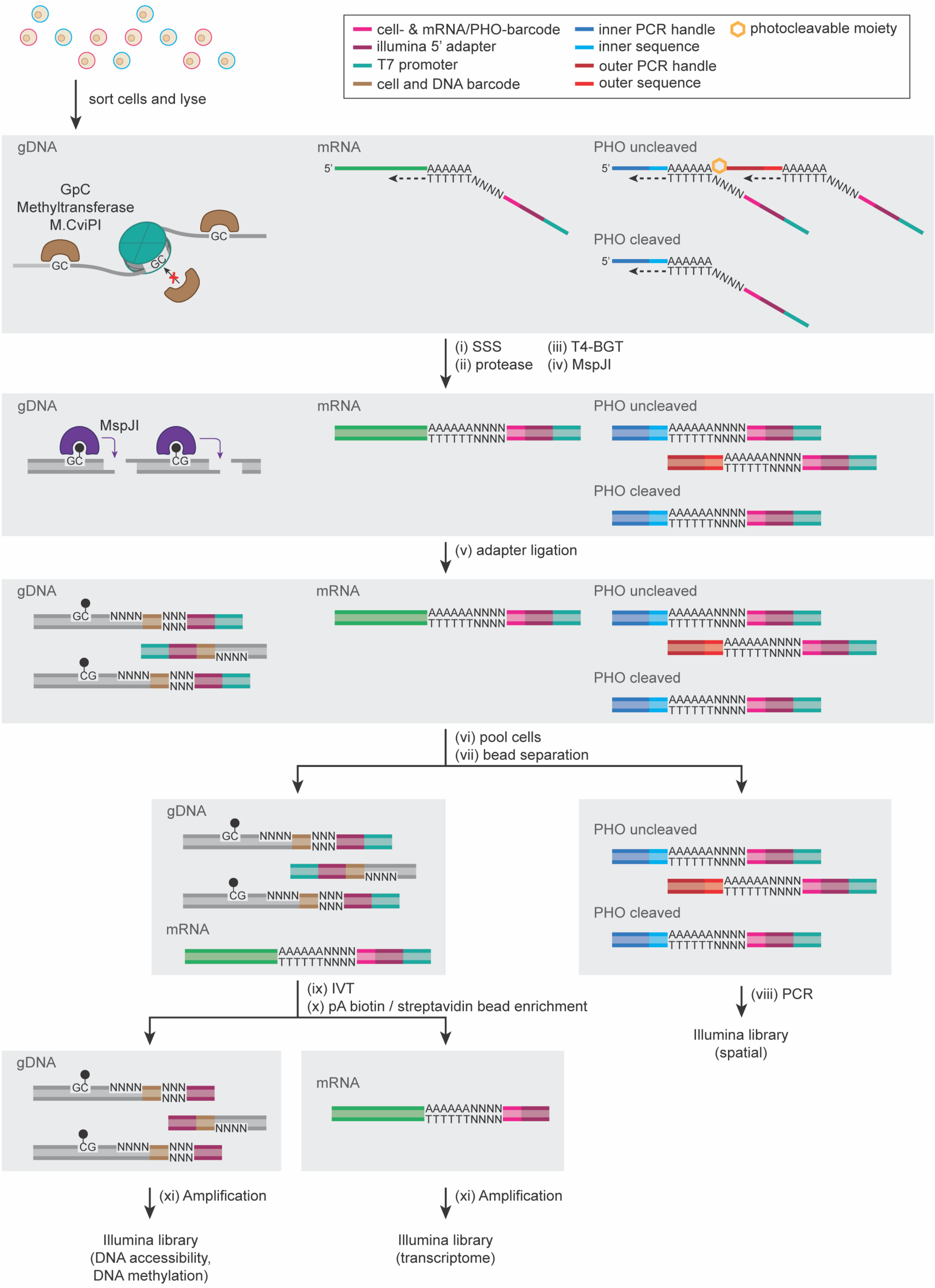

cell- & mRNA/PHO-barcode
illumina 5' adapter
T7 promoter
cell and DNA barcode
inner PCR handle
inner sequence
outer PCR handle
outer sequence
photocleavable moiety
sort cells and lyse
gDNA
GpC Methyltransferase M.CviPI
GC
mRNA
PHO uncleaved
PHO cleaved
AAAAAA
TTTTTTNNNN
(i) SSS
(ii) protease
(iii) T4-BGT
(iv) MspJI
MspJI
CG
AAAAAANNNN
TTTTTTNNNN
(v) adapter ligation
NNNN
NNN
(vi) pool cells
(vii) bead separation
(ix) IVT
(x) pA biotin / streptavidin bead enrichment
(viii) PCR
Illumina library (spatial)
(xi) Amplification
Illumina library (DNA accessibility, DNA methylation)
Illumina library (transcriptome)

**Supplementary Figure 12 | scSTAMP-MAT-seq library preparation pipeline.** Schematic of scSTAMP-MAT-seq to spatially profile mRNA, DNA accessibility and DNA methylation from the same cell . First, cells are incubated with PHOs and illuminated with light patterns. Single cells are then dissociated, sorted into 384-well plates, and lysed. The following two steps are then performed simultaneously – (1) mRNA is reverse transcribed and the 'inner' and 'outer' DNA sequences of PHOs are copied using a poly-T primer with an overhang containing a cell- and mRNA/PHO-specific barcode, a UMI, a 5' Illumina adapter and a T7 promoter; and (2) a methyltransferase, M.CviPI, is used to methylate cytosines in a GpC context within open chromatin. Next, after second strand synthesis (i), gDNA chromatin is stripped using protease (ii), 5hmC sites in the genome are glucosylated to block downstream detection by the restriction enzyme MspJI (iii), and MspJI is subsequently added that recognizes methylated cytosines in the genome and creates double-stranded (ds)-DNA breaks (iv). The digested genomic DNA (gDNA) molecules are ligated to ds-adapters containing a cell- and gDNA-specific barcode, a UMI, the 5' Illumina adapter and a T7 promoter (v). Following this, as all molecules are tagged with cell- and molecule-of-origin-specific barcodes, individual wells of the 384-well plate are pooled (vi), and the short PHO-derived molecules are separated from barcoded endogenous mRNA- and gDNA-derived molecules using SPRI beads (vii). PHO-derived molecules are then amplified by PCR to generate spatial Illumina libraries (viii). The mRNA- and gDNA-derived molecules are amplified by IVT (ix), followed by a further enrichment using polyA biotin/streptavidin coated beads to separate mRNA-derived molecules from gDNA-derived molecules (x). Finally, the remaining mRNA- and gDNA-derived molecules are amplified (xi) to generate endogenous transcriptome and epigenome Illumina libraries (see Methods for details).

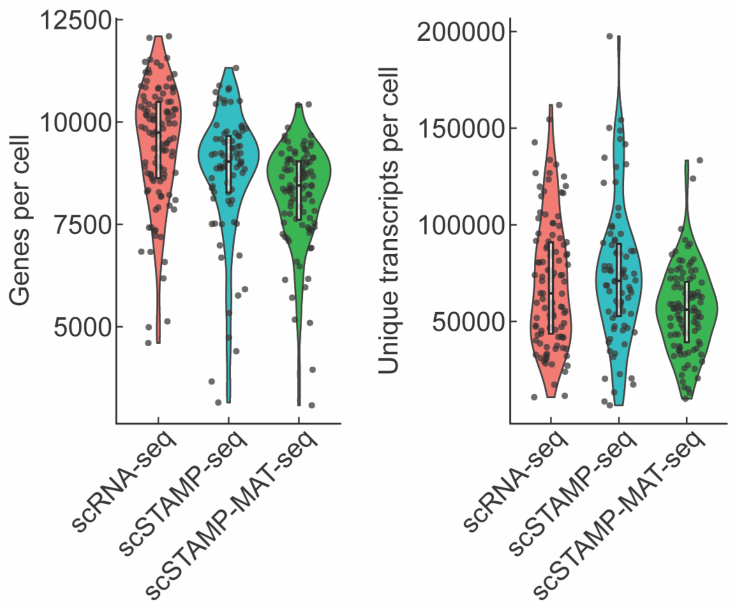


**Supplementary Figure 13 | Comparison of the transcriptome obtained from scSTAMP-MAT-seq to scRNA-seq and scSTAMP-seq.** Violin plots of the number of genes and total number of unique transcripts detected per cell for standard scRNA-seq, scSTAMP-seq and scSTAMP-MAT-seq.

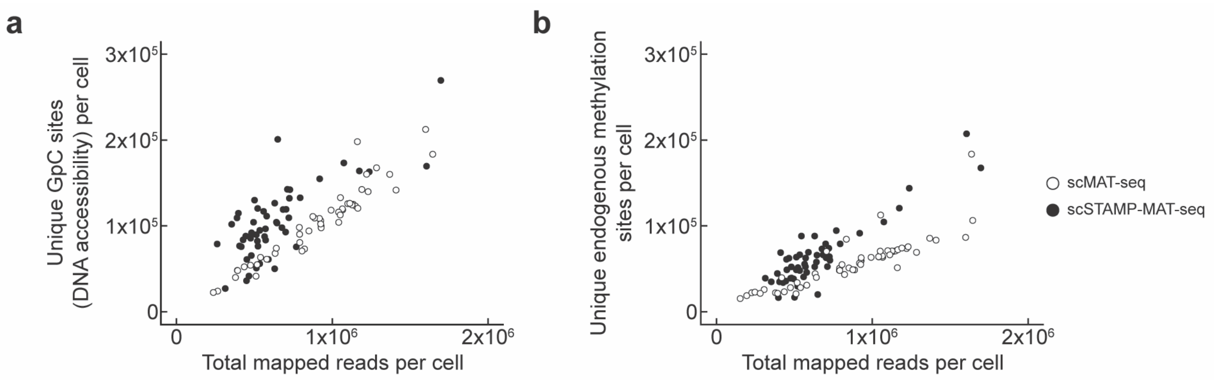


**Supplementary Figure 14 | Benchmarking the epigenome of scSTAMP-MAT-seq to scMAT-seq.** **a**, Scatterplot of unique methylated GpC sites (DNA accessibility) detected per cell as a function of the sequencing depth for scSTAMP-MAT-seq and MAT-seq. The number of unique methylated GpC sites detected per cell for scMAT-seq ranged from 40,000 to 212,000, with a median of 109,000. The number of unique methylated GpC sites detected per cell for scSTAMP-MAT-seq ranged from 27,000 to 269,000, with a median of 96,000. **b**, Scatterplot of unique endogenously methylated CpG sites detected per cell as a function of the sequencing depth for scSTAMP-MAT-seq and scMAT-seq. The number of unique endogenous methylation sites detected per cell for scMAT-seq ranged from 21,000 to 106,000, with a median of 59,000. The number of unique endogenous methylation sites detected per cell for scSTAMP-MAT-seq ranged from 16,000 to 207,000, with a median of 55,000.
